# Spatial Decoding of Tertiary Lymphoid Structure Maturation in Non-Small Cell Lung Cancer Using Deep Neural Networks

**DOI:** 10.64898/2026.01.16.697961

**Authors:** Bob Chen, Conrad Foo, Alma Andersson, Brandon D. Kayser, Yiming Yang, Alsu Missarova, Priyanka Kulkarni, Jan-Christian Huetter, Sharmila Chatterjee, Youngmi Kim, Yan Liang, Emily Killinbeck, Sarah Murphy, Eloisa Fuentes, Jennifer M. Giltnane, Raj Jesudason, Bo Li, Aicha BenTaieb, David Richmond, Tommaso Biancalani, Robert J. Johnston, Eric Lubeck, Tyler Risom, Orit Rozenblatt-Rosen, Runmin Wei, Lisa M. McGinnis

## Abstract

Understanding the role of tertiary lymphoid structures (TLS) is crucial in non-small cell lung cancer (NSCLC), as they are associated with patient prognosis and treatment outcomes. Specific cellular ecosystems that originate anti-tumor activity or predict immunotherapy response remain poorly characterized. To this end, we developed a high-resolution, multimodal spatial atlas jointly profiling transcriptomics, proteomics, and histology to characterize TLS maturation in NSCLC alongside secondary lymph organs as a baseline. Using this atlas, we proposed a pathologist-in-the-loop framework that combines a variational graph autoencoder (VGAE) with diffusion pseudotime to refine human expert annotations and characterize TLS maturation. These spatial molecular representations were extended to H&E whole-slide images via a vision transformer-based foundation model. Next, we resolved cellular composition, spatial organization, and cell-cell interactions within these data and defined two divergent spatial ecosystems. Clinical evidence suggests that these ecosystems are associated with distinct patient outcomes: a mature germinal center niche with favorable prognoses, and a tumor-macrophage-fibroblast niche with unfavorable prognoses. In summary, our work decodes key components of TLS heterogeneity, identifies hallmark spatial patterns involved in NSCLC adaptive immunity, and provides a framework for translating spatial omics insights into clinical applications.

## Main

Non-small cell lung cancer (NSCLC) represents approximately 85% of lung cancer diagnoses, contributing substantially to global cancer mortality. Despite significant histologic and molecular heterogeneity within this disease, NSCLC is broadly classified into adenocarcinoma (LuAD) and squamous cell carcinoma (LuSC) histologies ^1,2^. A subset of NSCLC tumors contain an immune-rich tumor microenvironment (TME) composed of diverse innate and adaptive immune cells. Immune checkpoint blockade (ICB) therapies, such as those targeting the PD-1/PD-L1 axis, modulate these cell populations and have demonstrated a survival benefit in responders ^3^. Previous studies have highlighted key components of the TME associated with robust humoral responses and anti-tumor immunity ^4,5^.

Tertiary lymphoid structures (TLSs) are highly organized aggregates of immune cells that form ectopically in non-lymphoid tissues ^6^. TLSs evolve from immature lymphoid aggregates (LAs), characterized by loosely clustered lymphoid cells, with minimal organization, to mature TLS (TLSm), with well-defined B cell follicles, germinal centers (GC) and T-cell zones (TZ) ^6^. Although TLS presence and maturity often correlate with favorable prognosis or immunotherapy response, the mechanisms by which TLS mediate antitumor immunity remain incompletely characterized^5,6^. B lymphocytes such as plasma cells have been increasingly implicated in local humoral responses and antigen presentation functions, but their precise roles in distinct TLS ecosystems are not yet resolved ^4,5^. To understand their clinical associations, we constructed a multimodal spatial atlas of NSCLC TLS and adjacent TMEs. Leveraging deep learning models on high-dimensional spatial and imaging data, we analyzed cellular composition, spatial architecture, histopathology, and cell-cell interactions to map the functional heterogeneity of TLS ecosystems.

This multimodal spatial atlas integrates single-nucleus RNA-seq with high-resolution spatial transcriptomics (CosMx SMI), proteomics (MIBIscope), and histopathology (haematoxylin and eosin; H&E), and spans 180 fields of view (FOVs) from 8 patients, capturing the TLS developmental spectrum alongside secondary lymphoid organ (SLO) follicles. To analyze these data, we developed a geometric deep learning framework based on a variational graph autoencoder (VGAE), to jointly model spatial organization and cellular transcription complemented by label transfer from paired snRNA-seq. We further applied a pretrained digital pathology foundation model to transfer molecular insights into histopathological contexts. This integrated analysis enabled the comparison of functional niches across the NSCLC TME, TLS, and SLOs, revealing novel spatial signatures of TLS maturation and critical TME interactions. Notably, the prognostic relevance of these signatures was confirmed in large patient cohorts (TCGA) and clinical trial data (OAK) ^7,8^. Finally, we demonstrated that these molecular insights can be directly applied to H&E data, establishing a scalable framework for clinical translation.

## Results

### Multi-modal Spatial Profiling Highlights Regional Heterogeneity within the NSCLC Tumor Microenvironment

We collected 8 NSCLC and 2 tonsil tissue blocks for spatial and single-nucleus (sNuc) profiling (Fig. 1a). These tissue blocks were sectioned serially with slices designated for H&E, CosMx, and MIBIscope. For the CosMx assay, a 30-gene panel was designed to target the TME in addition to the baseline 1000-gene panel. Similarly, a 41-plex panel was designed for protein targeting with MIBIscope (Supplementary Table 1). Additional slices were used to generate a paired sNuc reference atlas using the Chromium Flex whole transcriptome platform. Each serially sectioned modality was registered to the same coordinate system using WSIReg ^9^, enabling the annotation of transcripts by H&E. LA and TLS annotations were dependent on the observation of compact, ovoid collections of lymphocytes with diameter constraints. TLSm were identified by the observation of well-formed germinal centers (GCs), contrasting TLSi, or immature TLS, which were associated with either ill-defined GCs or high endothelial venule (HEV) densities. These histological designations were made assuming that LAs may represent diverse cellular communities preceding the formation of primary and secondary follicle-like structures, lacking the supportive stroma to further organize follicle zonation or maturation ^6,10^. Tumor nests were identified as dense regions of cells with hyperchromatic and pleomorphic nuclei surrounded by stromal desmoplasia. Aligning with CosMx FOVs, the following regions-of-interest (ROI) were annotated: tumor nests (n = 68), lymphoid aggregates (LA, n = 40), immature tertiary lymphoid structures (TLSi, n = 7), mature tertiary lymphoid structures (TLSm, n = 9), and lymphoid follicles (LF, n = 78) in tonsil samples. Detailed annotation strategies and specimen metadata are described in Methods section and Supplementary Table 2, Extended Data Fig. 1a.

**Figure 1:**
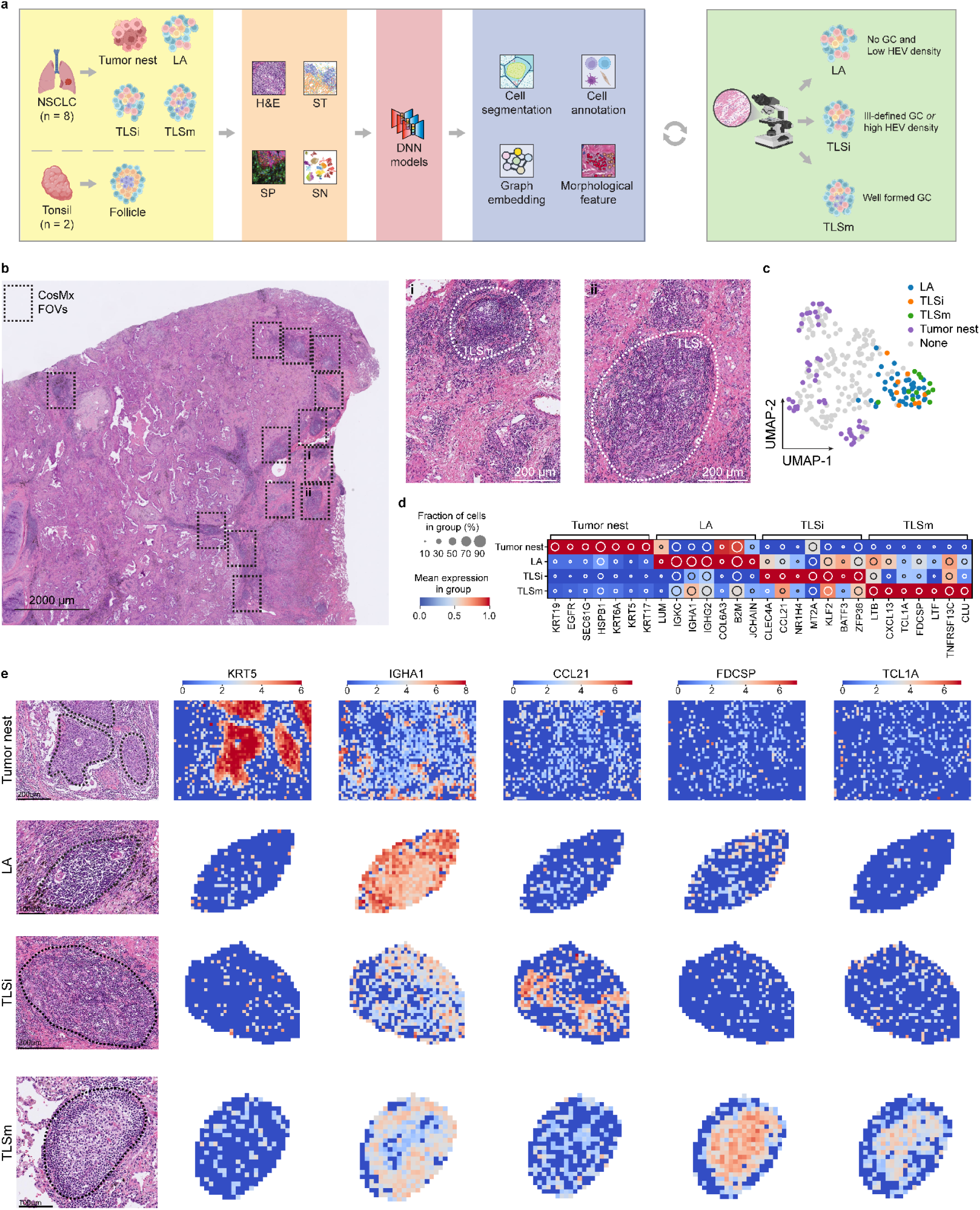
Multi-modal Spatial Profiling Highlights Regional Heterogeneity within the NSCLC Tumor Microenvironment. **(A)** Overview of multi-modal spatial data collection and analysis starting from sample setup focusing on ROI, alignment with H&E, spatial transcriptomics, spatial proteomics, and sNuc transcriptomics. These data are then processed with respective methodologies relevant for cell segmentation, annotation, spatial graph embedding, and morphological feature learning. FOVs and ROI are selected by histopathological analysis. **(B)** H&E of an NSCLC tissue sample, scale bar = 2000µm. Outlines FOVs assayed with CosMx with highlights containing TLSm (i) and TLSi (ii), scale bar = 200µm. **(C)** UMAP of pseudobulked transcripts per ROI, annotated by histological classification **(D)** Dotplot depicting ROI-based differential gene expression by histological classification, values scaled after log normalization. **(E)** Segmentation-free visualization of key differentially expressed genes *KRT5, IGHA1, CCL21, FDCSP*, and *TCL1A* within spatially-binned grids, with each FOV divided into 2,500 bins of about 258µm^2^ each. Transcripts were log normalized per spatial bin. For tumor nest and TLSi FOVs, scale bar = 200µm; for LA and TLSm, scale bar = 100µm.

Leveraging deep neural network (DNN) models across these paired modalities, we performed a series of spatial analyses including: 3D cell segmentation on the DAPI with CellPose followed by Baysor segmentation, multi-level cell type annotations (L1 for cell type, L2 for cell subtype) on the spatial transcriptomics (ST) data with scVI and scANVI, spatial niche analysis with a VGAE model, and H&E-based deep learning of TLS maturation with CellViT ^11–15^. We highlighted a pathologist-in-the-loop modeling paradigm to enhance our histopathological annotations with DNN-based ROI identification (Fig. 1a).

Based on the annotated ROI (Fig. 1b, Extended Data Fig. 1b), we performed aggregated, pseudobulk gene expression analysis using the ST data, which revealed substantial tumor heterogeneity. In contrast, LA and TLS exhibited higher transcriptional similarity (Fig. 1c, Extended Data Fig. 1c), consistent with their observed proteomic characterizations using MIBIscope (Extended Data Fig. 1d). Next, we conducted segmentation-free analysis by spatially binning the detected transcripts into a square grid. Differential gene expression (DEG) analysis identified top marker genes for distinct ROI and associated cell types and programs (Fig. 1d, Supplementary Table 3). In tumor nests, keratin genes along with *EGFR*, were highly expressed. Overall, LAs, TLSi, and TLSm showed an enrichment in B cell related genes such as *TNFSF13* (Fig. 1d). In LAs, immunoglobulin-related (*IGHG2, IGHA1*) genes were also enriched, consistent with plasma cells and their mucosal IgA-positive variants. Additionally, mesenchymal genes (*LUM, COL6A3*) were detected, reflecting early fibroblast associations with LA; in contrast, chemokines such as *CCL21* were upregulated in TLSi. Extending this, TLSm displayed robust *LTB, CXCL13, TCL1A, FDCSP*, and *LTF* expression, representing concentrated lymphoid and mesenchymal cell communities. To further explore differences between TLS and SLO, we included two tonsil samples in a corresponding spatial analysis and observed an enrichment of genes we had found in the TLSm within tonsil LF, such as *FDCSP* and *TCL1A* (Extended Data Fig. 1e,f).

In addition, these genes also varied spatially (Fig. 1e, Extended Data Fig. 2a-e). For example, plasma cell markers like *IGHA1* were expressed interspersed within LAs but also across the peripheral boundaries of keratin-enriched tumor nests. In contrast to the diffuse *CCL21* expression in TLS, we observed tonsil LFs enriched for *CCL21* at the interface of their adjacent interfollicular regions. This chemokine was enriched in TLSi and previously characterized as a facilitator of T cell recruitment through interactions with *CCR7* ^10^. As expected, TLS maturation markers *FDCSP* and *TCL1A* were focally expressed in the germinal centers (GC) of TLSm and tonsil LF. This is consistent with previously described components of functional maturation such as germinal center B cells (GC-B) and follicular dendritic cells (FDC), further supporting their role in TLS development and immune organization. Importantly, the spatial enrichment of these transcriptional programs supports the H&E-anchored annotation of lung TLS alongside their comparison across tissue contexts, such as with tonsil LFs.

### Mature TLS in NSCLC Share Cellular Hallmarks of Secondary Lymphoid Organ Germinal Centers

To address ST imaging challenges associated with high cell density and overlapping cells in the z-dimension, we applied a two-step segmentation pipeline leveraging Cellpose ^12^ and Baysor ^11^. This approach improved segmentation accuracy by integrating 3D DAPI-staining with the 3D spatial distribution of transcripts (Extended Data Fig. 3).

Subsequently, we employed the DNN frameworks, scVI and scANVI ^13,16^, to transfer cell labels from the paired sNuc reference data (Fig. 2a). This reference atlas was finely annotated using a similar strategy with annotations derived from Salcher et al. ^17^. To address the potential platform effects between single-cell reference and spatial query data sets, as well as mitigating the stochasticity of DNN models, we devised an ensemble scVI and scANVI voting pipeline to leverage the joint latent space of CosMx and sNucRNA-seq data for robust label transfer.

**Figure 2:**
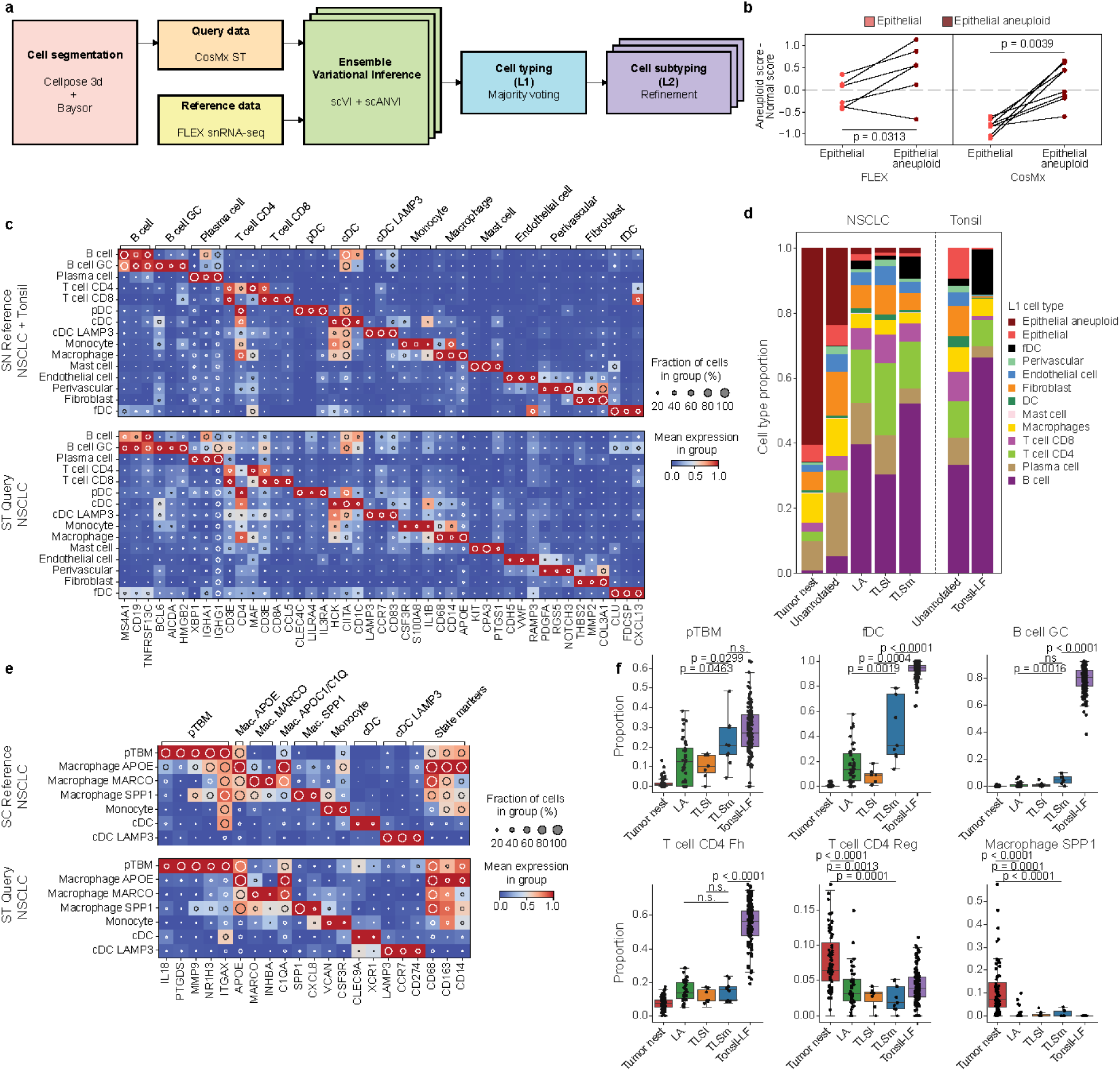
Mature TLS in NSCLC Share Cellular Hallmarks of Secondary Lymphoid Organ Germinal Centers. **(A)** Schematic of CosMx imaging data processing, from transcript location and DAPI staining to annotated, high-resolution cell subtypes through the application of Cellpose 3d, Baysor, and hierarchical co-embedding of paired snRNA-seq data through scVI and scANVI. **(B)** Sample-wise evaluation of dichotomizing normal and aneuploid epithelial cells using one-sided Wilcoxon signed-rank tests for mean values across spatial and sNuc modalities. **(C)** Dotplot depicting relevant high-resolution cell subtypes across the SN reference, derived from NSCLC and Tonsil samples, and NSCLC ST query dataset; values scaled after log normalization. **(D)** Stacked barplot showing proportional representation of L1 cell types captured across all classified ROI. **(E)** Dotplot depicting high-resolution myeloid cell subtypes within NSCLC-derived SN and ST datasets, highlighting *IL18, PTGDS, MMP9, NR1H3*, and *ITGAX* expression in pTBMs; values scaled after log normalization. **(F)** Boxplots depicting L2 cell subtype representation, with proportions normalized by cell lineage per ROI. Mann-Whitney U tests were performed between ROI classifications per subtype.

To differentiate aneuploid epithelial from normal epithelial cells in NSCLC samples, we applied CopyKat ^18^ to infer copy number profiles from the paired sNuc reference data and derived a signature score. Briefly, the underlying genes of this score were calculated through the DEGs of predicted aneuploid and diploid epithelial cells (Methods) ^19^. We found significant increases of this score in aneuploid epithelial cells compared to diploid cells through a paired, one-sided Wilcoxon signed-rank test in both sNuc and CosMx modalities (Methods, Fig. 2b, Supplementary Table 4). Next, more broadly across cell types, we performed DEG analysis and observed consistent top DEGs expression patterns between SC reference and ST query datasets, both in the context of our NSCLC ST and tonsil ST data (Fig. 2c, Extended Data Fig. 4a, Supplementary Table 5,6). The consistency between data modalities was further demonstrated by Spearman correlation analyses across cell types (Extended Data Fig. 4b,c, Supplementary Table 7).

Using the H&E-derived annotations across our NSCLC and Tonsil samples, we compared the distribution of different L1 and L2 cell subtypes (Fig. 2d, Extended Data Fig. 4d). This showed an expected distribution of epithelial and stromal cell presence that skewed to tumor nests. Within these tumor nests, a diversity of immune cells was also present, including macrophages, T cells, and B/plasma cells. In contrast, B and T lymphocytes were more enriched in the LA, TLSi, TLSm, and tonsil. Interestingly, LA, TLSi, and TLSm looked similar in L1 cell type frequency. TLSm show FDC and B cell trends indicative of more advanced maturation like that seen in tonsil LFs. In these LFs, we also observed a decrease in stromal cell representation (fibroblasts and endothelial cells) and CD8^+^ T cells compared to immune aggregates in NSCLC, reflecting structural and functional differences between tertiary and secondary lymphoid structures. We further characterized the detected myeloid cell populations, due to our observation of CD68^+^ cells within TLSm using MIBI, identifying seven distinct myeloid subtypes, with their marker gene expression profiles shown in Fig. 2e and Extended Data Fig. 4e. These states included putative tingible body macrophages (pTBM), which express *IL18*, *PTGDS*, *MMP9* and are *CD68* positive (Extended Data Fig. 4e,f). Additionally we identified CD14^+^ monocytes alongside tumor associated macrophage populations (TAMs) differentiated by *SPP1*, *MARCO*, or *APOE* expression. We also identified conventional dendritic cells (cDCs), which express *CLEC9A* and *XCR1*, and LAMP3^+^ mature cDCs, each exhibiting distinct expression patterns of immune-related genes. These diverse myeloid cell subtypes highlight the functional heterogeneity within the NSCLC microenvironment, with specific subpopulations likely contributing to immune modulation and TLS maturation.

When comparing ROI by lineage-normalized L2 cell subtype composition, we observed an increasing trend in pTBM, FDCs, and germinal center B (GC-B) cells during TLS maturation, with their highest enrichment observed in tonsil LFs (Fig. 2f). In the B lymphocyte compartment, additional contrasts include IGHA^+^ plasma cells, with LAs containing proportionally more than TLSm, suggestive of a mucosal or extrafollicular plasma cell population (Extended Data Fig. 4d). Mesenchymal-lineage cells such as fibroblastic reticular cells (FRCs), expressing *CCL19* and *CXCL12*, were depleted within the cores of these LFs relative to LA and TLSi. In the T lymphocyte compartment, CD4^+^ T follicular helper (Tfh) cells were enriched in tonsil LFs, reinforcing their role in GC formation and B cell activation. In contrast, CD8^+^ exhausted T cells (Tex), CD4^+^ regulatory T cells (Tregs) and SPP1^+^ macrophages were predominantly detected in tumor nests, suggesting their involvement in immunosuppressive interactions within the tumor microenvironment. Finally, CD3^+^ T cells expressing Naive-like markers such as *SELL*, *TCF7*, and *CCR7* were depleted in tumor nests relative to LAs and TLS. We demonstrate that by applying integrative methods we enable the joint utilization of information across spatial and sNuc transcriptomic modalities for pinpointing critical cell subtypes at high resolutions.

### A Pathologist-in-the-loop Modeling Framework Integrates Transcriptomics and Histology for Characterizing TLS Maturation

We constructed a VGAE model to integrate gene expression profiles and cellular spatial locations from CosMx ST data (Methods), capturing a spatially-aware latent representation of cellular organization within NSCLC tissue blocks (Fig. 3a). UMAP embeddings derived from the VGAE latent space (**z**) revealed a pseudotime trajectory (PT) that corresponded to TLS maturation (Fig. 3b). Cells with higher VGAE-PT values were predominantly located within germinal center regions of TLS, indicating that the pseudotime captures key features of TLS organization and maturation. To investigate how pseudotime progression varies across different cell subtypes, we compared their cumulative distribution function (CDF) of PT values. Using the Kolmogorov-Smirnov (K-S) test, we identified significant differences in PT distributions across cell populations (Fig. 3c, Supplementary Table 8). Notably, GC B cells and FDCs exhibited the highest PT values, suggesting their association with advanced stages of TLS maturation. In contrast, naive B cells, memory B cells, and pTBM exhibited intermediate PT values, suggesting a transitional state in the maturation continuum, whereas other cell types displayed significantly lower VGAE-PT values. The K-S statistics heatmap further highlights the degree of separation between cell subtypes, reinforcing the structured progression of cellular differentiation and maturation within the spatial niche. A comprehensive VGAE-PT distribution and K-S statistics on all cell subtypes are included in Extended Data Fig. 5a.

**Figure 3:**
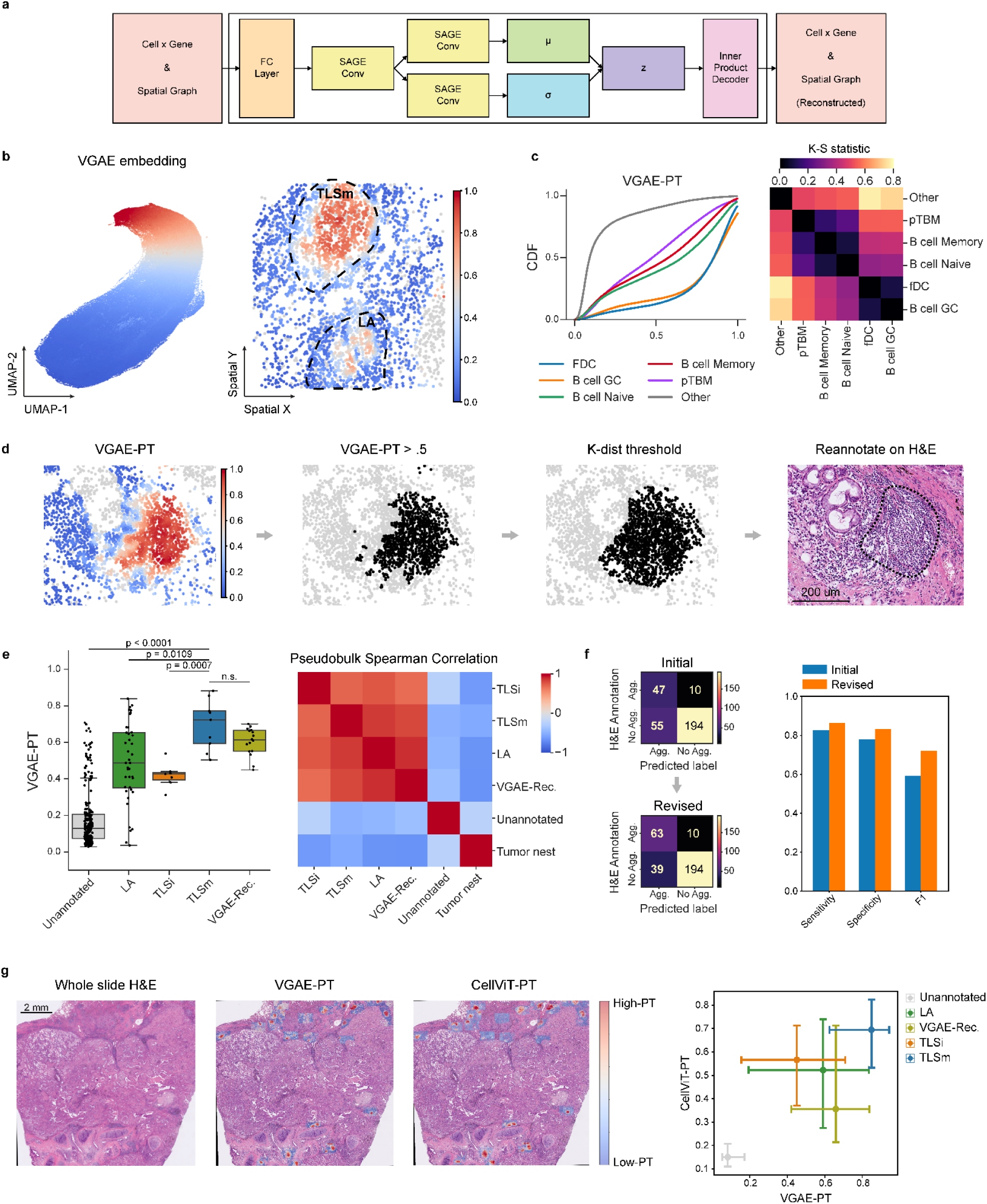
A Pathologist-in-the-loop Modeling Framework Integrates Transcriptomics and Histology for Characterizing TLS Maturation. **(A)** Schematic of VGAE architecture. Initial steps encode the input transcriptional and spatial information through a FC layer, followed by parallel SAGE convolutional layers producing a latent representation which is ultimately learned through reconstructing the original spatial gene expression and graph. **(B)** UMAP of VGAE latent space with VGAE-PT overlaid (left). Corresponding spatial representation demonstrates enrichment of high pseudotime or spatial maturation values within annotated ROI such as TLSm and LA (right) **(C)** CDF of VGAE-PT values partitioned by L2 cell subtype, with only the top 5 cell types highlighted: FDCs, B cell GC, B cell Naive, B cell Memory, and pTBMs (left). Corresponding K-S test statistic shown for each curve, where asterisks indicate p < 0.05, full matrix in Supplementary Table 7 (right). **(D)** Protocol for the de novo detection of candidate ROI for pathologist-in-the-loop feedback; starting with VGAE-PT thresholding at 0.5 and defining polygonal regions based on K-distance and region size thresholding, scale bar = 200µm. **(E)** Boxplot of mean VGAE-PT values per ROI partitioned by histopathological annotation or VGAE-Rec. status. Mann-Whitney U tests were performed between ROI classifications (left). Pairwise Spearman correlations of pseudobulked ROI highlighting VGAE-Rec. transcriptional similarity to initial ROI identified (right). **(F)** Initial and revised confusion matrices of ROI classifications identifying aggregate versus no aggregate, given VGAE-Rec. candidate ROI (left) alongside sensitivity, specificity, and F1 diagnostic metrics (right). **(G)** Visualization of whole slide H&E with overlays showing VGAE-PT values per FOV and CellViT-PT, with high-PT representing more mature LA/TLS (left) alongside a scatterplot comparing CellViT-PT and VGAE-PT values, 25% – 75% quantiles were added (right).

Notably, we identified ROI with high VGAE-PT values that were not fully annotated by pathologists in the initial round of annotation. To systematically refine the annotation of these potential immune aggregates, we applied spatial cutoffs on high scoring ROI (Methods, Fig. 3d). The identified ROI were then subjected to a second round of pathologist annotation using corresponding H&E-stained tissue sections, leading to identification of additional TLS-like ROI. These ROI were then classified as “VGAE recovered regions” (VGAE-Rec.).

Then, we examined pseudotime differences across annotated ROI, finding that TLSms demonstrated higher PT values than TLSi and LA, while unannotated ROI showed lower values (Fig. 3e). Interestingly, LA displayed a broader range of PT values than TLSi and TLSm, likely reflecting more diffuse spatial organization compared to the highly structured TLS. As expected, the recovered ROI also demonstrated high VGAE-PT values. Spearman correlation of pseudobulked gene expression and L2 cell subtype composition confirmed these VGAE-Rec. ROI are more similar to TLS/LA than tumor nests and other unannotated ROI (Fig. 3e, Extended Data Fig. 5b). Of the initial 55 VGAE-identified, TLS-like ROI, 16 of them were validated by pathologists. This iterative process identified additional immune aggregates and improved our prediction accuracy by increasing the F1 scores, sensitivity and specificity (Fig. 3f). These results underscore the value of integration of ST-based DNN models like VGAE with expert pathological annotation to better identify and characterize ROI in complex tissue architectures.

To further extend the applicability of our DNN-based framework, we leveraged VGAE-PT as a reference and applied a pretrained imaging foundation model, CellVit ^15^, to paired whole-slide H&E images. We trained a two-layer prediction model on the deep feature vector extracted by CellVit to predict PT values across the tissue (Fig. 3g, Extended Data Fig. 5c). The model successfully captured key ST patterns, with TLSm exhibiting the highest predicted PT values and unannotated ROI showing the lowest, consistent with expectations. While the Spearman correlation between CellViT-PT and VGAE-PT values was 0.48 in training and 0.38 in testing, the model effectively distinguished ROI with different maturation statuses. These findings highlight that deep features extracted from H&E images align with transcriptomics-derived pseudotime dynamics, demonstrating the potential of pretrained imaging models for large-scale spatial analysis.

Summarizing this synergistic, pathologist-in-the-loop modeling framework: the human-annotated data serves as an initial ground truth, enabling the ST-based VGAE model to recover additional, pathologist-validated ROI, while the imaging foundation model, guided by the learned pseudotime, expands spatial analysis to entire tissue sections (Extended Data Fig. 5d,6). By combining transcriptomic and histological data with human expertise, this framework provides a scalable and powerful solution for uncovering spatially organized regions in complex tissue environments. Together, these findings show how learned spatial trajectories can uncover latent tissue organization, enhance expert annotation, and extend spatial insights across whole-slide images.

### A Spatial Pseudotime-derived, Multi-cellular Signature of TLS Maturation Predicts Clinical Benefit from ICB

We next developed a novel gene signature, termed the *VGAE score*, derived from a logistic regression model with L1 penalty predicting high VGAE-PT versus low VGAE-PT ^20^(Supplementary Table 9, methods). In addition, we compared this score to previously identified signatures which predict Atezolizumab response in NSCLC: a broad signature derived from patients with improved OS after Atezolizumab treatment (*ICB response score*) and an intratumoral plasma cell signature (*Plasma cell score*) ^4^. When examining the composition of individual ROI, we found that the ICB response score targeted both intratumoral plasma cells as well as mature TLS (Fig. 4a, Extended Data Fig. 7a). The Plasma cell score was elevated in stromal areas, often near tumor nests, whereas the VGAE score was more restricted to organized lymphoid structures (Fig. 4b). We also observed correlations between MHCII-related gene expression to both the ICB response and our VGAE score (Extended Data Fig. 7b). This dichotomy was further supported by K-dist analysis showing plasma cells were located nearer to tumor nests, while B cells were predominantly near TLS and LF (Fig. 4c).

**Figure 4:**
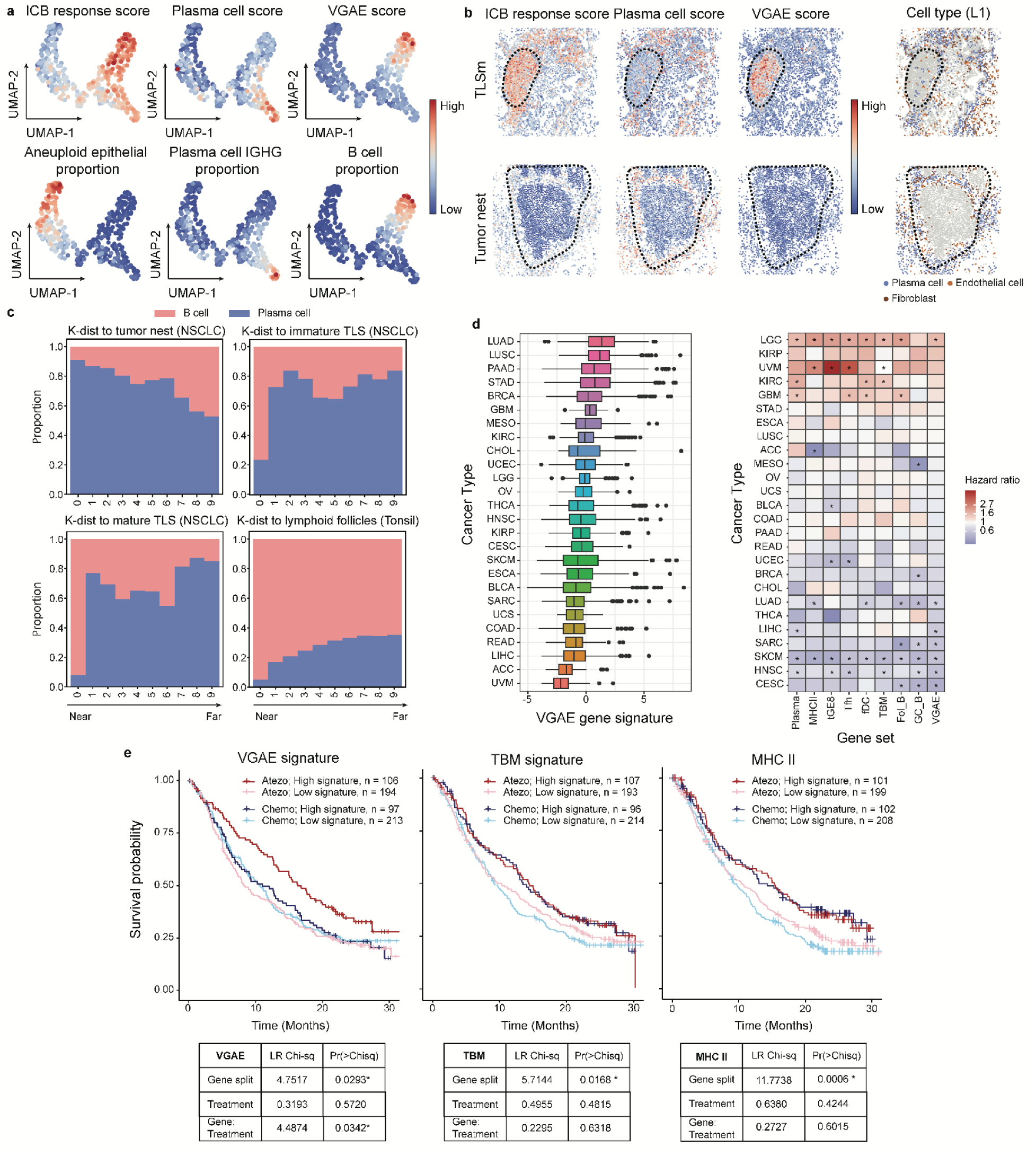
A Spatial Pseudotime-derived, Multi-cellular Signature of TLS Maturation Predicts Clinical Benefit from ICB. **(A)** UMAP of ROI by L2 cell subtype composition demonstrating gene signature score representation within ROI alongside cell subtype proportions **(B)** Spatial visualizations of gene signature scores in relation to key ROI exemplified by a TLSm (upper) and tumor nest (lower), key stromal and lymphoid cell populations are visualized (right) **(C)** Stacked barplot showing proportional representation of B cells and Plasma cells across tissue contexts, distributed into 10 K-dist bins starting from near or within the ROI. **(D)** Boxplot of GSVA scores calculated for VGAE gene signature across TCGA cancer types (left). Overall survival and hazard ratios given GSVA gene set scores quantizing patient groups, asterisks indicate p < 0.05 **(E)** Kaplan-Meier plots for the VGAE, TBM, and MHC II gene signatures scored by GSVA in the OAK cohort, dichotomization procedure detailed in methods. Tables contain ANOVA results with 1 degree of freedom, full results in Supplementary Table 10.

To assess broader relevance, we evaluated the VGAE score across 27 TCGA cancer types using GSVA^21^. This score was significantly enriched in cancers known for immune interactions (LUAD, LUSC, and PAAD) and depleted in immune-cold or privileged tumors (UVM, and ACC) (Fig. 4d) ^22–25^. We then investigated the prognostic value of these spatially-derived signatures in TCGA with respect to overall survival. The VGAE score and related, previously defined immune signatures (i.e. GC_B, Fol_B, etc.) generally showed favorable survival in immune-infiltrated cancers such as LUAD, SKCM, and HNSC but paradoxically correlated with worse outcomes in immune-privileged or excluded types such as UVM, LGG, and KIRC (Fig. 4d) ^4,26^. These findings underscore that the prognostic significance of TME components is tied to their spatial organization and overall immune context of the cancer type.

Finally, we examined the association of these signatures with ICB outcomes using data from the OAK phase III NSCLC trial (Atezolizumab vs. Docetaxel)^8^. Multivariate Cox proportional hazard analysis revealed significant prognostic differences when splitting the cohort by the VGAE, TBM and MHC II signatures, respectively. More importantly, we also observed an interaction between VGAE scores with Atezolizumab treatment (ANOVA, Gene:Treatment p = 0.0342, Supplementary Table 10, Fig. 4e). Modeling each treatment arm independently showed that patients with high VGAE scores had improved overall survival in the Atezolizumab arm; this improvement was not observed in the chemo arm (HR; atezolizumab high versus low HR = 0.65 [0.48–0.87], p = 0.0003; chemo high versus low HR = 1.00 [0.75–1.32], p = 0.9760). The MHCII signature score was prognostically beneficial, associated with improved overall patient survival regardless of treatment arm and had no significant Gene:Treatment interaction (Fig. 4e, Methods, ANOVA, Gene:Treatment p = 0.6015, atezolizumab high versus low HR = 0.74 [0.55–1.00], p = 0.0484; chemo high versus low HR = 0.66 [0.49–0.89], p = 0.0058). A similar trend was observed in correlated signatures targeting pTBMs and tGE8s (Extended Data Fig. 7c,d). These signatures were compared to the Plasma cell signature, which showed a significant, Atezolizumab-specific enhancement of efficacy in a subset of patients (ANOVA, Gene score:Treatment p=0.0235; atezolizumab high versus low HR = 0.59 [0.43–0.79], p = 0.0004; chemo high versus low HR = 0.94 [0.70–1.24], p = 0.6470). Collectively, these analyses demonstrate that spatially informed signatures, identified by our VGAE, capture distinct TLS-specific niches which hold significant impact on improved survival across cancers and provide insights into TME configurations relevant to immunotherapy response.

### Spatial Analysis Reveals Cellular Ecosystems with Divergent Axes of Adaptive Immunity in NSCLC

To investigate the spatial distribution of each cell type relative to different annotated ROI, we calculated the spatial K-dist to tumor nests, immature and mature TLS in NSCLC, and LF in tonsils (Fig. 5a, Extended Data Fig. 8a). Our analysis revealed that SPP1^+^ TAMs, along with FAP^+^ cancer-associated fibroblasts (CAFs), CD8^+^ exhausted T cells (CD8 Tex), regulatory T cells (Tregs), and cytotoxic T cells were near tumor nests. Conversely, FDCs and germinal center B cells (GC B) were near both immature and mature TLS, along with naive and memory B cells. Notably, in TLS, naive-like and cycling T cells were nearer relative to cytotoxic, exhausted, and regulatory T cells. Near tonsil LFs, we observed FDCs and GC B cells with pTBM, CD4^+^ T follicular helper cells (Tfh), and naive B cells, whereas naive-like T cells were more distant (Fig. 5a,b, Extended Data Fig. 8a).

**Figure 5:**
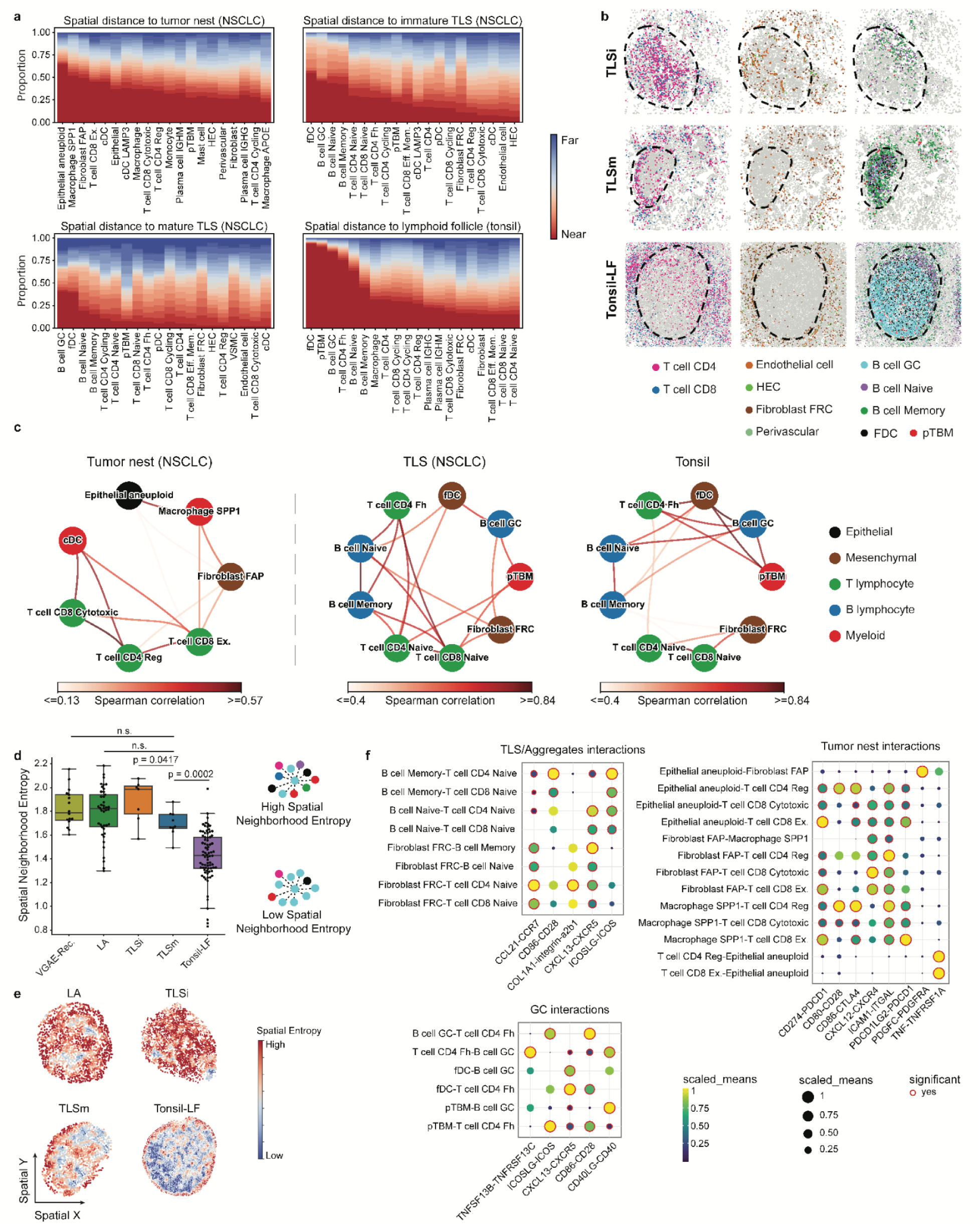
Spatial Analysis Reveals Cellular Ecosystems with Divergent Axes of Adaptive Immunity in NSCLC. **(A)** Stacked barplots visualizing L2 cell subtype representation across 20 K-dist bins calculated on the basis of each individual tissue block. Cells within the boundaries of ROI were placed in the nearest bin, shown in red. **(B)** Spatial plots annotated by ROI per row and subtypes per column. ROI were determined by H&E registration. Left column shows CD4 and CD8 T cells; middle column shows Endothelial cells, HECs, FRCs, and Perivascular cells; right column shows GC, Naive, and Memory B cell alongside other follicle-associated cell subtypes such as FDCs and pTBMs **(C)** Graph visualizing top 3 cell subtype co-localizations through Spearman correlation of K-dist values. Nodes annotated by cell lineage; edges colored by Spearman correlation values between connected nodes **(D)** Boxplot of mean spatial neighborhood entropy metric across ROI partitioned by histopathological annotation. Mann-Whitney U tests were performed between ROI classifications (left). Example of cell neighborhoods with high and low entropy, where colored nodes represent unique cell subtypes and dotted edges represent neighborhood adjacency (right). **(E)** Single-cell spatial neighborhood entropy values overlaid on core neighborhood nodes across representative ROI **(F)** CellPhoneDBv5-predicted ligand-receptor interactions within spatially-defined microenvironments, representing likely signaling occurring within TLS/Aggregates, Tumor nests, and GCs.

We then calculated pairwise Spearman correlations on the K-dist matrix between each L2-level cell subtype to assess cell type colocalization patterns (Extended Data Fig. 8b). We highlight two distinct colocalization communities using hierarchical clustering (Fig. 5c). The NSCLC tumor nest community was enriched with aneuploid epithelial cells, SPP1^+^ TAMs, FAP^+^ CAFs, Tregs, Texs, and cDCs, suggesting an immunosuppressive microenvironment. The colocalization of SPP1^+^ TAMs and FAP^+^ CAFs indicates potential roles in immune evasion and stromal remodeling, while the presence of Tregs and exhausted T cells highlights a dysfunctional T cell landscape within the tumor nest. In contrast, the NSCLC-TLS community showed a strong colocalization among B and T lymphocytes, along with pTBMs, supportive FRCs, and FDCs.

Notably, we observed that naive-like CD4^+^ and CD8^+^ T cells, naive and memory B cells, and T follicular helper (Tfh) cells were closely colocalized in NSCLC-TLS, consistent with early TLS formation and immune activation. In tonsil tissues, a germinal center-enriched subcommunity of FDCs, GC-B cells, pTBMs, and Tfh was strongly colocalized, reflecting mature germinal center structures of SLO. To further assess the spatial differences between LA/TLS and SLO, we devised a structural entropy metric where lower values indicate high levels of cell subtype spatial partitioning and organization^27^ (Methods). Here, we observed a significant decrease in mature TLS compared to immature TLS, followed by tonsil LF with the lowest entropy. (Fig. 5d,e). This indicates an increased spatial organization during TLS maturation, though this organization did not reach the same level observed in tonsil LF. Interestingly, this entropy was relatively diverse in LAs, indicating that histologically-identified LAs may contain a wider range of molecular variation than detectable through H&E, which was consistent with our VGAE-PT observations (Fig. 3e). To identify cell-cell interactions within various spatial compartments, we performed CellPhoneDB analysis after restricting cell types into three spatial microenvironments: tumor nest-enriched, TLS/Aggregate-enriched, and GC-enriched (Fig. 5f). Within the tumor nest, enriched pathways involving PD-L1/PD-L2-PD-1 alongside CD86-CTLA4 suggest immune checkpoint interactions originating from tumor cells^28^. Concurrently, Treg-related signaling via CD80-CD28 was observed originating from macrophages and tumor cells^29^. Interactions facilitating inflammation-related adhesion through the ICAM1-ITGAL axis between tumor cells, fibroblasts, macrophages, and T cells were also observed ^30,31^. Further, the detection of CXCL12-CXCR4 and TNF-TNFRSF1A interactions pointed towards angiogenic signaling alongside a localized inflammatory or tumor necrosis response ^32,33^.

Contrasting tumor nests, the TLS/Aggregate-enriched community showed upregulated interaction networks characteristic of immune recruitment and the development of adaptive immunity. Interactions crucial for lymphocyte homing, such as the CCL21-CCR7 and CXCL13-CXCR5 axes, were evident. Corroborated by localized cell type enrichments, these were potentially mediated by FRCs and other stromal cells, alongside structural interactions involving the collagen-integrin axis ^34^. Within these developing structures, pathways promoting lymphocyte activation were also evident. T cell costimulation, mediated by CD86-CD28 and ICOSLG-ICOS interactions, were observed between memory B cells and naive-like T cells ^35,36^. These same co-stimulatory pathways were further enhanced in more mature, GC-enriched microenvironments, particularly between GC B cells and Tfh, highlighting their potential role in sustaining Tfh function and B cell affinity maturation ^37,38^. Furthermore, essential signals for robust B cell responses within GCs were detected, such as CD40-CD40LG interactions, vital for B cell activation, class switching, and memory formation ^39^. Supporting this spatially-restricted adaptive response, we also observed BAFF receptor (TNFSF13B-TNFRSF13C) signaling, crucial for B cell survival ^39,40^.

In parallel, we defined another transcriptional signature reflecting the immunosuppressive tumor niches defined in our spatial data, characterized by FAP^+^ CAFs and SPP1^+^ TAMs (*the FAP-SPP1 score*), which was also enriched in cancers like LUAD and LUSC (Extended Data Fig. 8c). Conversely, the FAP-SPP1 score, marking specific tumor-associated stromal and macrophage niches, demonstrated a consistent unfavorable impact across diverse cancers, particularly strong in immune-privileged settings (UVM, LGG, and KIRP), suggesting a potential role in pro-tumorigenic immune evasion. Importantly, the FAP-SPP1 signature was associated with significantly worse survival, demonstrating a potentially negative prognostic role (Extended Data Fig. 8d; ANOVA, Gene score p = 0.0243, Supplementary Table 10).

Our spatial analysis in NSCLC and tonsil tissues revealed distinct cell type distributions and spatial organizations across tumor nests, TLS, and SLO. Tumor nests were enriched with aneuploid epithelial cells and immunosuppressive interactions, while TLS and germinal centers exhibited structured immune cell colocalizations, with increased B and T cell activation signals. By integrating spatial organization metrics and cell-cell interaction analysis, we detected key molecular interactions involved with TLS maturation, highlighting fundamental differences between NSCLC-TLS and SLO. These results also demonstrate that TLS and tumor nests in NSCLC are shaped by orthogonal spatial patterns and signaling programs, pointing towards spatially-resolved mechanisms that may inform immunotherapy strategies.

## Discussion

We introduce a single-cell resolution spatial atlas of TLS in NSCLC, integrating multiple data modalities, including single-nucleus RNA-sequencing, spatial transcriptomics, spatial proteomics, and H&E imaging, through deep learning frameworks to uncover the cellular and spatial heterogeneity of these structures. Our approach reveals complex spatial dynamics underlying TLS maturation and identifies cellular niches with prognostic and therapeutic relevance. Crucially, we compared TLS architecture to well-characterized SLOs, and delineated both preserved lymphoid features (e.g., follicular zonation, stromal scaffolding) alongside molecular deviations unique to TLS in the NSCLC TME. In parallel, we contrasted TLS-specific immune interactions with those of tumor nests, highlighting differences between supportive and suppressive microenvironments.

Our computational framework was accelerated through the application of deep learning on several fronts. First, we improved cellular spatial resolution using 3D segmentation and bayesian approaches, consistent with recent findings across multiple spatial platforms ^45^. Second, we devised a consensus voting procedure to ensure robust, hierarchical cell type label transfer. Third, we applied trajectory inference to our spatially-aware VGAE latent space to establish a TLS maturation pseudotime. Fourth, by integrating H&E imaging foundation models with this VGAE model, we enabled the prediction of ST-based features directly from imaging data. This end-to-end approach not only revealed fine-grained cellular and spatial dynamics of TLS, but also allowed molecular insights to be transferred directly to H&E images, supporting scalable clinical translation.

Our VGAE score, associated with ICB response in OAK, represents a robust FDC presence: from CXCL13-mediated B cell recruitment to supporting GC maturation. This aligns with recent findings by Patil et al., implicating plasma cells as key mediators of TLS-related ICB outcomes^41–44^. Interestingly, we observed that some LAs contained increased IGHA^+^ plasma cells relative to TLSm, suggesting extrafollicular plasma cell differentiation potentially primed by tumor or CAF-mediated TGF-B signaling ^45^. We also found that TLSi did not consistently represent transitional states between LAs and TLSm. Although TLSi display features of immune recruitment and stromal support, some of them may lack key elements for full maturation, suggesting they represent a more complex niche rather than a simple transitional stage between LAs and TLSm.

The incorporation of SLO spatial transcriptomics alongside TLS allowed for the identification of human pTBMs. Their conserved localization across SLO and TLS, suggests a shared role in active germinal centers, likely phagocytosing apoptotic B cells that had undergone somatic hypermutation (SHM) ^46,47^. In addition, previous investigations of TBM functioning suggest that they may be sources of B cell-modulating prostaglandins in the germinal center, a transcriptional pattern corroborated by our observation of pTBMs specifically expressing *PTGDS* ^48^. Interestingly, prostaglandins have also been implicated in an ICB response-predictive signature in melanoma with TLS ^44^. While these pTBMs appear generally associated with TLS GCs, the wider relationship between their potential SHM-responsive role and the B cell affinity maturation process in TLS remains unknown. On the other hand, myeloid colocalizations unique to tumor nests, particularly SPP1^+^ TAMs, contribute to immunosuppressive interactions. Similar colocalizations have been reported in CRC ^49^, where they restrict T cell infiltration and decrease cytotoxic responses. Consistent with these findings, we observed that this TAM-rich NSCLC tumor nest niche is correlated with worse clinical outcomes, suggesting immune suppression and pro-tumorigenic support.

Emerging evidence points towards additional factors influencing TLS-mediated anti-tumoral immunity. For instance, hypoxic TME conditions or tumor tryptophan metabolism surrounding TLS, in NSCLC and HCC, may influence PD-1 blockade response ^50,51^. Furthermore, the limitations of dissecting these complex 3D structures using primarily 2D spatial transcriptomics should be acknowledged ^52^. While this atlas spans a number of FOVs and multimodal contexts, the limited sample number may restrict the generalizability of our findings. Overall, the present work offers a diverse, single-cell resolution spatial atlas of TLS within the NSCLC TME. We decode this spectrum of maturation, yielding spatial signatures with clinical relevance alongside potential cell-cell interactions mediating this outcome, enabling further exploration into TLS-centric immunotherapeutic development.

## Methods

### CosMx SMI assay and data generation

FFPE blocks were serially sectioned using a microtome to a thickness of 5 μm for both NSCLC and Tonsil samples, and subsequently processed using Nanostring’s CosMx SMI Semi-Automated Slide Preparation for RNA Assays protocol (Nanostring, MAN-10159-02). Stained sections served as a guide for FOV selection, where 20 FOVs per block were selected using H&E annotations to capture a diversity of microstructures, with an enrichment for LA, TLS, and GCs. Sections were mounted on the back of the slide labeling area of a Superfrost Plus Micro Slide (VWR, 48311-703). These samples were then processed with the CosMx SMI platform with appropriate flow cell settings with respect to source tissue, using the 1000-Plex Human Universal Cell Characterization RNA Panel with 30 additional custom genes. For morphology visualization, B2M/CD298, PanCK, CD20, CD3 antibodies and DAPI were used for IF. These data were transferred to Nanostring’s AtoMx platform for downstream analysis and reprocessing.

### Flex single-nucleus assay and data generation

Two to three tissue sections, each between 25 and 50 µm thick, were initially deparaffinized. This was achieved by washing each section three times with 3 mL of xylene for 10 minutes. After deparaffinization, the tissue sections were rehydrated through ethanol baths. Each bath comprised 1 mL of ethanol and lasted 30 seconds: the sections were treated two times with 100% ethanol, followed by treatments with 70% ethanol, 50% ethanol, and finally water.

After rehydration, the tissue sections were subjected to two additional washes with Buffer U and Buffer V from Miltenyi’s FFPE Tissue Dissociation Kit (cat #3130-118-052). Subsequently, the sections underwent further digestion using the dissociation mix provided in the kit. Tissue disruption was performed using an Octodissociator in Tube C for 45-50 minutes.

10X Chromium Fixed RNA Profiling gene expression assay was performed as per manufacturer’s instruction (user guide-CG000527, kits-cat # 1000422, #1000547). Libraries were sequenced on an Illumina NovaSeq with paired-end dual-indexing (#1000251).

### Flex single-nucleus transcriptome processing

All the sc/snFFPE data were processed via the 10x Cell Ranger Flex Gene Expression pipeline. Specifically, tonsil samples were processed using Cell Ranger v7.0.1 with Chromium Human Transcriptome Probe Set v1.0; lung samples (all the rest) were processed using Cell Ranger v7.1.0 with Chromium Human Transcriptome Probe Set v1.0.1. The Transcriptome reference GRCh38 (GENCODE v32/Ensembl98) was used for all samples.

### Segmentation and transcript assignment

Each CosMX FOV (10 x 5472 x 3648 pixels, with 0.75 microns per z-pixel and 0.18 microns per x/y-pixel) was segmented independently. The FOV image was segmented using cellpose (fine-tuned cyto2 model) in 3D mode on the DAPI channel to generate 3D nuclear masks ^12^. Transcripts assigned to the nuclear masks were then used as the prior segmentation to the Baysor algorithm ^11^, with a prior confidence of 0.8. The scale parameter was estimated using the size model of cellpose, using the average radius across all z-stacks.

### CosMx single-cell quality control

An initial filtering step was performed removing cells with low confidence transcript assignments, using the baysor_avg_assignment_confidence metric output by default. Next, all CosMx datasets underwent cell filtering based on library size and unique gene detection thresholds. These metrics were calculated and log transformed with scanpy’s calculate_qc_metrics and the median absolute deviations (MAD) were calculated for each. Per sample, upper and lower bounds of 1 MAD were set for both metrics, yielding a set of candidate cells per metric. These intersecting cells were filtered out, likely representing missegmented cells with artifactual transcript assignments.

### Ensemble variational inference and label transfer

Label transfer was performed using our matched and annotated tonsil and NSCLC Flex sNuc datasets as reference. We devised an ensemble label transfer pipeline which was performed on the intersection of genes in the CosMx panel and the Flex transcriptome. This ensemble method involves 2 major phases: ensemble scANVI and post-processed majority voting. Due to discrepancies between the Flex and CosMx modalities alongside the inherent stochasticity of variational inference, we performed 10 replicate runs of scVI followed by scANVI. scVI parameters were as follows: n_latent = 32; max_epochs_scvi = 100; dropout rate = 0.3; and batch_key = “Block”, where each individual sample was classified as a separate block. scANVI parameters were as follows: max_epochs_scanvi = 100; n_samples_per_label = (100, 300, 500), where each value is run with its own set of 10 replicates.

This resulted in 30 distinct runs of scANVI classification, and each of these were post-processed with a classification entropy filter followed by consensus voting. Since each replicate has an associated cell ID by cell type classification probability matrix, the cell type classification entropy was calculated cell-wise and thresholded by finding the lowest local minimum by examining the histogram, at 0.4. Cells with classification entropies above this threshold are annotated as “Unknowns”, interpreted as mixed single-cell transcriptomes that could not be confidently assigned to a single cell type. Finally, consensus voting is performed and the final annotation is assigned by majority vote given a proportional threshold of .5, where classifications below this number are again assigned as unknowns. These ensemble runs were implemented as a Papermill workflow on compute nodes using Nvidia Tesla V100 32GB GPUs.

This scANVI-based joint embedding approach was also applied to cell lineage-specific subsets of both the reference and CosMx data, facilitating high-resolution L2 cell subtype annotation. High quality embeddings were selected from the ensemble by calculating the average silhouette score for separating reference cell types. A pathologist-in-the-loop strategy was then implemented, involving iterative evaluation of marker gene expression concordance between single-cell and spatial data to refine cell labels.

### Spatial k-nearest neighbor distance (k-dist)

Spatial kNN-distance (k-dist), were computed as the average distance from each cell of interest to its k-nearest neighbors of a specified target category, whether a specific cell type or an annotated spatial structure such as lymphoid aggregates. Using the Scikit-learn NearestNeighbors function, these distances are calculated for each cell per individual block. For further visualizing and comparison, the k-dists were divided into 20 bins using *qcut* from *pandas*. Cells within the borders of annotated ROI comprised the first bin.

### Spatial compositional analysis

Compositional analyses were conducted at two levels of cell type resolution (i.e., L1 and L2) across spatial contexts. L1 cell type counts were normalized per spatial context with the total cell number within that grouping as the denominator, thus the stacked barplots representing L1 cell types normalize to the total number of cells found within an ROI classification such as LA, TLSm, etc. L2 cell subtype analyses, depicted as boxplots, were normalized using their respective L0 (cell lineage-level) total cell numbers per spatial context as the denominator due to sparser L2 cell counts. For example, CD4 T follicular helper cells were normalized to the total number of T lymphoid cells found within individual ROI as opposed to the total cells across all lineages.

### Spatial kNN compositional entropy

We calculated spatial entropy based on spatial kNN graphs for each cell. First, we computed the spatial kNN within each tissue block using Squidpy’s spatial_neighbors function with coord_type=’generic’ and n_neighs=20. Next, for each cell, we calculated the Shannon entropy of its spatial neighbors on L2 cell type annotations using the entropy function from scipy.stats. In addition, we also calculated a mean value of spatial entropy for each annotated ROI.

### Co-localization analysis

To perform spatial co-localization analysis, we first computed the spatial k-dist from each single cell to every L2 cell type. These distances were then z-scored within each tissue block. This produced an *n x m* spatial distance matrix, where n is the number of cells and m is the number of L2 cell types. From this matrix, we calculated an *m x m* Spearman correlation matrix to assess co-localization patterns at the L2 cell type level. We applied hierarchical clustering with complete linkage on euclidean distances as implemented in scipy.cluster to organize L2 cell types into co-localization modules for downstream analysis. For graph visualization, we used Netgraph library to visualize modules, showing the top three edges per node based on Spearman correlation coefficients.

### CopyKat analysis and aneuploid signature scoring

CopyKAT 1.1.0 was applied to the NSCLC Flex count matrix to infer the copy number aberrations and identify aneuploid epithelial cells with following parameters: ngene.chr=5, win.size=25, KS.cut=0.05, distance= “euclidean”. We then performed differential gene expression analysis using scanpy rank_genes_groups with a minimum log2 fold change of 1. This test was done between predicted aneuploid and diploid epithelial cells pseudobulking these categories per sample. Top differentially expressed genes were then ranked and filtered by logFC, p-value, and mean expression. Cells were then scored using scanpy score_genes and the difference between these scores yielded the final score across the Flex and CosMx datasets.

### CellphoneDB receptor-ligand predictions with microenvironments

Using the preprocessed and phenotyped CosMx data, we first subset a maximum of 2,000 cells for each L2 cell type. Then, we performed CellphoneDB 5.0.1 ligand-receptor prediction using “statistical_analysis” with following parameters: iterations = 100, threshold = 0.05, result_precision = 3, pvalue = 0.05. We constructed a microenvironment data based on our co-localization analysis results: the tumor nest niche contains diploid and aneuploid epithelial cells, T cell CD8 Ex., T cell 8 Cytotoxic, T cell CD4 Reg, Macrophage SPP1, Fibroblast FAP and cDC; the TLS niche contains Fibroblast FRC, B cell Memory, B cell Naive, T cell CD4 Naive, T cell CD8 Naive; the GC niche contains T cell CD4 Fh, B cell GC, pTBM, and fDC; the Connective tissue niche contains Endothelial cell, Plasma cell IGHG, Plasma cell IGHA, Fibroblast, Plasma cell IGHM, VSMC, Fibroblast FN1, and HEC; and other cell types to the other group.

### Kaplan-Meier survival analyses

Survival analyses were performed using the signatures detailed in Supplementary Table 2 on both the OAK bulk RNA-seq dataset as described by Patil et al. and the TCGA pan-cancer bulk RNA-seq dataset ^4,7^. For the OAK dataset, genes were first filtered based on non-expression, then the gene sets were scored using GSVA using the ‘zscore’ method on the respective TPM log2 values. To understand the relationship between these gene sets, their pairwise Pearson correlations were calculated. Finally, each gene set was quantized using a tertile split and dichotomized by grouping the mid and low groups together to compare with the high group. Multivariate Cox proportional hazard (CoxPH) analyses were then performed using the *coxph* function examining survival as a function of the gene split,treatment arm and the interactions between these two. We then performed ANOVA test on these fitted models to understand the contributions of these factors to survival. For TCGA pan cancer analysis, we applied a similar approach, but without the inclusion of treatment arms as these were not a part of the TCGA dataset.

For this analysis we focused on solid tumors, so we excluded the control samples (CNTL), pilots (FPPP), and following cancers due to confounding effects: Lymphoid Neoplasm Diffuse Large B-cell Lymphoma (DLBC, hematologic malignancy), Kidney Chromophobe (KICH, sparse endpoints, use is not recommended), Pheochromocytoma and Paraganglioma (PCPG, small sample size, use is not recommended), Testicular Germ Cell Tumors (TGCT, germ cell malignancy, sparse endpoints, use is not recommended), Prostate adenocarcinoma (PRAD, hormone-driven malignancy, sparse endpoints, caution advised), and Thymoma (THYM, sparse endpoints, use is not recommended) ^53^.

### Deep morphological feature extraction

Serial H&E sections were aligned to the CosMx data using WSIReg ^9^ with both rigid and nonlinear B-spline transforms. The FOV images were composited into a whole slide image based on the global CosMx coordinates, and the mutual information loss was computed only in the valid FOV image regions. A vision transformer-based model, CellViT-SAM-H ^15^, was used to extract deep morphological features from the aligned serial sections in a tile-wise fashion, with a tile size of 1024×1024 and a 64 pixel overlap. This generated a 1280-dimensional deep feature vector for each 16×16 spatial patch. Morphological image features were then mapped to CosMx cells by taking the patch that contained the cell centroid.

### Morphological deep feature analysis

Morphological features for each cell were used as regressors to predict the VGAE-PT of cells that were excluded from transcriptomics analysis. A simple two layer neural network was trained to predict the VGAE-PT from the extracted morphological feature vector for each cell. The first layer transformed the full feature vector into a 25-dimensional latent space, and the second layer predicted the pseudotime from the latent features. A sigmoid activation function was used in the hidden layer. The model was trained using gradient descent with MSE loss. The ADAM optimizer with a learning rate of 0.001 was used and the model was trained for 50 epochs; 20% of the cells were used for training and 80% of the cells were used for testing.

### Variational Graph Autoencoder (VGAE)

To complement traditional cell labeling approaches, we implemented a spatially aware method that integrates spatial context with molecular information. This was achieved using a modified Variational Graph Autoencoder (VGAE), originally introduced by Kipf and Welling, which extends the Variational Autoencoder (VAE) framework by incorporating graph-structured data.

VGAE is based on a probabilistic latent variable model, where an encoder maps input data to a structured latent space, and a decoder reconstructs relevant features from this representation. Given an input dataset *X* that represents each cell’s gene expression with an associated adjacency matrix *A* that details the cells’ spatial relationships we model the latent representation *Z* using an encoder network *f(.)*, which approximates the posterior distribution:

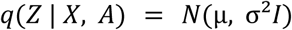

where the mean *μ* and variance *σ²* are learned through graph convolutional layers. The prior distribution on *Z* is assumed to be a standard normal distribution:

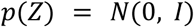

The model is trained by maximizing the evidence lower bound (ELBO), which consists of a reconstruction term and a regularization term:

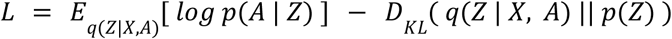

The decoder reconstructs the spatial adjacency matrix using an inner product decoder, which computes the probability of an edge between nodes *i* and *j* as:

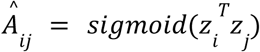

where *sigmoid(.)* denotes the sigmoid function. The reconstruction loss is formulated as a binary cross-entropy loss between the predicted adjacency matrix *Â* and the observed adjacency matrix *A*:

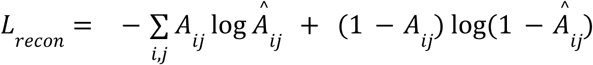

The variational regularization is imposed through the Kullback-Leibler (KL) divergence, ensuring that the learned distribution remains close to the prior:

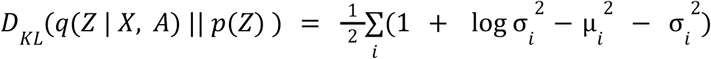

where summation is over all latent dimensions.

To enable gradient-based optimization, we use the reparameterization trick, rewriting the latent variable as:

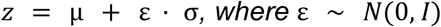

This allows gradients to propagate through the stochastic sampling step during backpropagation. The full objective function is optimized using stochastic gradient descent, with a weighting factor *β* applied to the KL divergence term, as in β-VAE formulations:

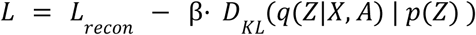

where β controls the trade-off between reconstruction fidelity and latent space regularization.

This approach enables the integration of spatial organization and molecular signatures into a unified latent representation, improving the accuracy of cell type annotation while preserving spatial relationships.

With respect to implementation, we next outline the exact architecture of our network and other relevant design choices, this deviates slightly from the original implementation. Given the gene expression input *X* and the spatial graph *A*, we do the following operations:

1. 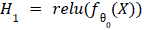
2. 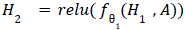
3. 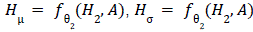
4. 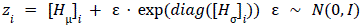
5. 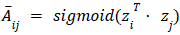

For computational efficiency, we replace the loss suggested in the original VGAE paper:

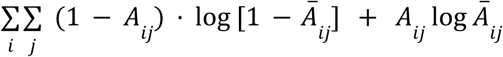

With a negative sampling strategy:

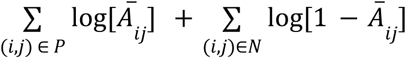

Where 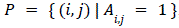, and *N* is a set of randomly sampled edges with the condition that 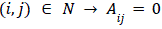 and |*N*| = |*P*|, *N* is resampled during each epoch. Furthermore, we let β = 1 / *M* where M is the number of nodes in the graph.

For our network layers 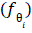, we let:

● 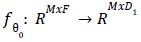, FC layer
● 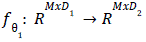, SAGEConv layer
● 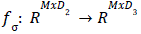, SAGEConv layer
● 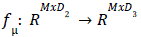, SAGEConv layer

Where 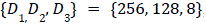, here *D*_3_ will be the size of the latent space.

The SAGEConv layer is defined as:

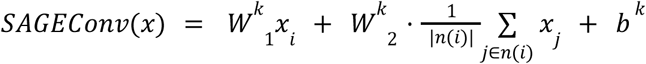

Where *n*(*i*) is the neighborhood of cell *i* and 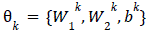 are learnable parameters for layer *k*. We implement our version of the VGAE in Python using PyTorch and PyG. All code with documentation is available in the associated code repository. We used PyTorch 2.3.1+cu121 and PyG 2.5.3, for a full specification of package versions see the code repository.

For training, we used the Adam optimizer, as implemented in the PyTorch suite with a learning rate of 0.001, (all other settings were set to default values). A batch size of 1 (one graph per batch) was used. We trained the model for 250 epochs. During training we apply a dropout function to the *Z* representations, where 50% of the values were randomly chosen and set to be zero.

Ultimately, graphs allow us to encode dependencies or relationships between observations, making them a well-suited choice for spatial data; where some dependency between proximal observations is usually expected.

In our case, we represent each FOV (field of view) as a graph, each cell is treated as a node connected to its K nearest (spatial) neighbors. The feature expression of each cell is assigned as attributes of the node representing that cell. Edges are represented as binary features, 1 indicating a connection between two nodes and 0 meaning no connection. In our analysis, we used *K* = 10.

### VGAE-PT calculation, guided spatial ROI recovery, and evaluation

In order to represent the immune microenvironment, we used a subset of the learned VGAE embedding focused on immune and stromal cells while excluding the tumor nests and adjacent ROI. On the VGAE latent space, we calculated the spatial pseudotime by diffusion map using sc.tl.diffmap, this was followed by performing diffusion pseudotime with 4 diffusion components, while rooting the pseudotime in a dense region of germinal center B cells ^54^.

Candidate VGAE recovered ROI were identified through a four-step process: 1. Pseudotime thresholding: Cells with VGAE-PT values above 0.5 were selected, indicating advanced pseudotemporal states consistent with more organized or mature lymphoid structures. 2. Spatial filtering: A k-dist (with k = 20) was computed, and cells within the top 20th percentile of spatial proximity were retained to define local regions. 3. Graph-based clustering: For each tissue block, a spatial neighbors graph was constructed using k equal to half the square root of the total number of detected cells. Leiden clustering was then performed at a resolution of 0.1. Spatial clusters with fewer than 100 cells or containing more than 10% epithelial cells were excluded.. 4 Pathological validation: The remaining ROI were mapped onto corresponding H&E images and reviewed by a pathologist following the same annotation protocol.

For precision-recall evaluation of this VGAE-PT workflow, all candidate ROI (VGAE-Leiden) identified were compared against the original ground truth (GT) annotated ROI. ROI-level label assignments were based on proportional overlaps: ROI with less than one-third overlap with GT annotations were labeled as false positives; GT ROI with less than one-third overlap with VGAE-Leiden clusters were labeled as false negatives; and ROI with more than one-third mutual overlap were considered true positives. Additionally, true negatives were defined as GT-negative areas where more than half the ROI showed no overlap with any VGAE-Leiden cluster. Within each field of view (FOV), unannotated areas were treated as a single negative sample.Candidate false positive and false negative ROI were then re-evaluated by a pathologist. Based on the second-round review, ROI labels were updated to reflect the revised ground truth. Validated VGAE-Leiden ROI were re-labeled as VGAE-Recovered, representing true positives that were previously missed in the first-round of annotation.

### VGAE signature score derivation with logistic regression

The underlying VGAE gene set used to calculate a signature score was derived from the mean feature coefficients of ten logistic regression rounds with a limited sampling procedure. Each run consisted of five primary steps, starting from the full CosMx dataset: 1) log normalization 2) sampling 0.5% of the total number of observations per L2 cell subtype 3) filtering low quality cells and nonspecific transcripts 4) feature selection through calculating the top 300 DEGs per downsampled L2 cell subtype using scanpy rank_genes_groups and taking the union 5) performing logistic regression using sklearn LogisticRegression to classify cells with VGAE-PT values greater than median TLSms value. This logistic regression step used Z-scaled values with an 80/20 train/test split using an L1 penalty with the inverse regularization strength C set to 3e-3 with a max iteration number of 500 and otherwise default parameters. Finally, the mean of these feature coefficients were calculated and sorted, followed by an additional filter removing non-lineage-specific gene expression. For example, if a gene was expressed in B lymphoid and myeloid lineage cells, it was removed from the signature due to nonspecificity.

### MIBIscope tissue preparation and imaging

Established protocols were followed for MIBIscope tissue processing and staining ^55^, antibody conjugation ^56^, and imaging ^57^. Briefly, tissue sections were baked at 70deg for 30 minutes, deparaffinized and rehydrated, and underwent heat induced epitope retrieval in a pH9 buffer (Agilent Dako, S2375). Sections were then blocked with a fish gelatin (Sigma-Aldrich G7765-250) and donkey serum (Sigma-Aldrich D9663-10ML) buffer, and stained overnight at 4deg with a cocktail of metal-conjugated antibodies in staining buffer ^58^. The following morning a secondary antibody solution is applied to the tissues, and slides are subsequently washed, fixed with glutaraldehyde (EMS, 16020), dehydrated, and stored in a nitrogen cabinet until imaging.

The panel of metal-conjugated antibodies is described in Supplementary Data File 1b. Metal-conjugated antibodies were either purchased directly from vendors (Ionpath, Standard Biotools) or generated following the Ionpath MIBItag protocol ^56^. Slides were imaged on the commercial MIBIscope platform at Ionpath HQ (Ionpath Inc, Menlo Park CA). For each pathologist-guided ROI, H&E topological markers were used to set MIBIscope FOVs centering around the structure. One 800×800um FOV was captured at 2048×2048 pixels on the “Fine” (∼550nm) imaging setting. ROI images were accessed using MIBItracker viewer (https://mibi-share.ionpath.com/). Image overlays for figures were generated on the MIBItracker viewer software.

## Data Availability

Data related to this manuscript will be deposited upon publication.

## Code Availability

Data related to this manuscript will be deposited upon publication.

## Acknowledgements

We are grateful for the organ and tissue donors and their families who participated in this study, OAK, and TCGA. Additionally we thank Mariya Barch, Barzin Nabet, and Akshay Krishnamurty for helpful discussions and feedback.

## Author contributions

LMM, ORR, and RJJ conceived the study. PK and SC performed all single-nucleus transcriptomics experiments. EK, SM, YK, and YL performed all spatial transcriptomics experiments. AA, AM, BC, BDK, CF, RW, and YY and analyzed all experiments. TR performed and analyzed spatial proteomics experiments. AA, BC, CF, and RW conceived of computational methods and applications. EF, JMG, and LMM performed all histopathological annotations. BC, CF, LMM, and RW wrote the manuscript and produced the figures. All of the authors edited the manuscript.

## Materials and Correspondence

Correspondence to Runmin Wei and Lisa M. McGinnis.

## Supplementary Table Information

**Supplementary Table 1: MIBIscope metal-conjugated antibodies and usage conditions**

MIBIscope metal conjugated antibodies used in this study are listed from lowest mass tag to highest. Metal tag, target protein, antibody clone, antibody vendor, catalog number, and stock concentration and titer are all included.

**Supplementary Table 2: NSCLC sample specimen characteristics and overall histopathological observations**

The overall histopathological annotations and clinical metadata are noted in this table for the eight FFPE NSCLC sample specimens used in this study; i.e. patient age, sex, disease staging, diagnoses, etc.

**Supplementary Table 3: Segmentation-free, ROI differential gene expression statistics**

Table related to Figure 1 and Extended Data Figure 1. Full list of differential gene expression statistics performed using scanpy rank_genes_groups in ‘wilcoxon’ mode between ROI classifications. Each tab separates two distinct sets of tests, which were run with or without the inclusion of Tumor Nests and Unannotated regions.

**Supplementary Table 4: Comprehensive list of gene signatures used in this study**

Table related to Figures 2,4 and Extended Data Figures 4,7, and 8. Includes gene set descriptions and originating sources alongside comprehensive gene sets.

**Supplementary Table 5: CosMx-detected cell subtype differential gene expression statistics**

Table related to Figure 2 and Extended Data Figure 4. Full list of differential gene expression statistics performed using scanpy rank_genes_groups in ‘wilcoxon’ mode between multiple cell subtype resolutions for CosMx data. Each tab separates distinct sets of tests, within lineage contexts and between lineage contexts across tissue sources.

**Supplementary Table 6: Flex-detected cell subtype differential gene expression statistics**

Table related to Figure 2 and Extended Data Figure 4. Full list of differential gene expression statistics performed using scanpy rank_genes_groups in ‘wilcoxon’ mode for Flex sNuc data. Each tab separates distinct sets of tests, within the myeloid lineage context and between lineage contexts.

**Supplementary Table 7: scANVI ensemble label transfer consistency and sNuc transcriptomic profile correlation**

Table related to Figure 2 and Extended Data Figure 4. Metrics related to scANVI ensemble label transfer consistency for annotating NSCLC and Tonsil CosMx data. Additionally contains diagnostic metrics calculated for the DEGs of these resultant annotations across Flex and CosMx modalities.

**Supplementary Table 8: K-S test statistic and significance matrices**

Table related to K-S tests in Figure 3 and Extended Data Figure 5. Full, pairwise K-S test statistics for visualized matrix plots.

**Supplementary Table 9: Gene coefficients across multiple logistic regression models identifying VGAE-PT-high cells**

Table containing full feature coefficient values of downsampled sampled logistic regression models. Tabs separate individual runs and final averaged values.

**Supplementary Table 10: OAK survival analyses and statistics**

Full CoxPH and ANOVA results which underlie Kaplan-Meier plots shown in Figure 4 and Extended Data Figure 8. Includes results from independently modeled treatment arms across each gene signature score tested.

## Extended Data Figures and Legends

Extended Data Figures and Legends

**Extended Data Figure 1:**
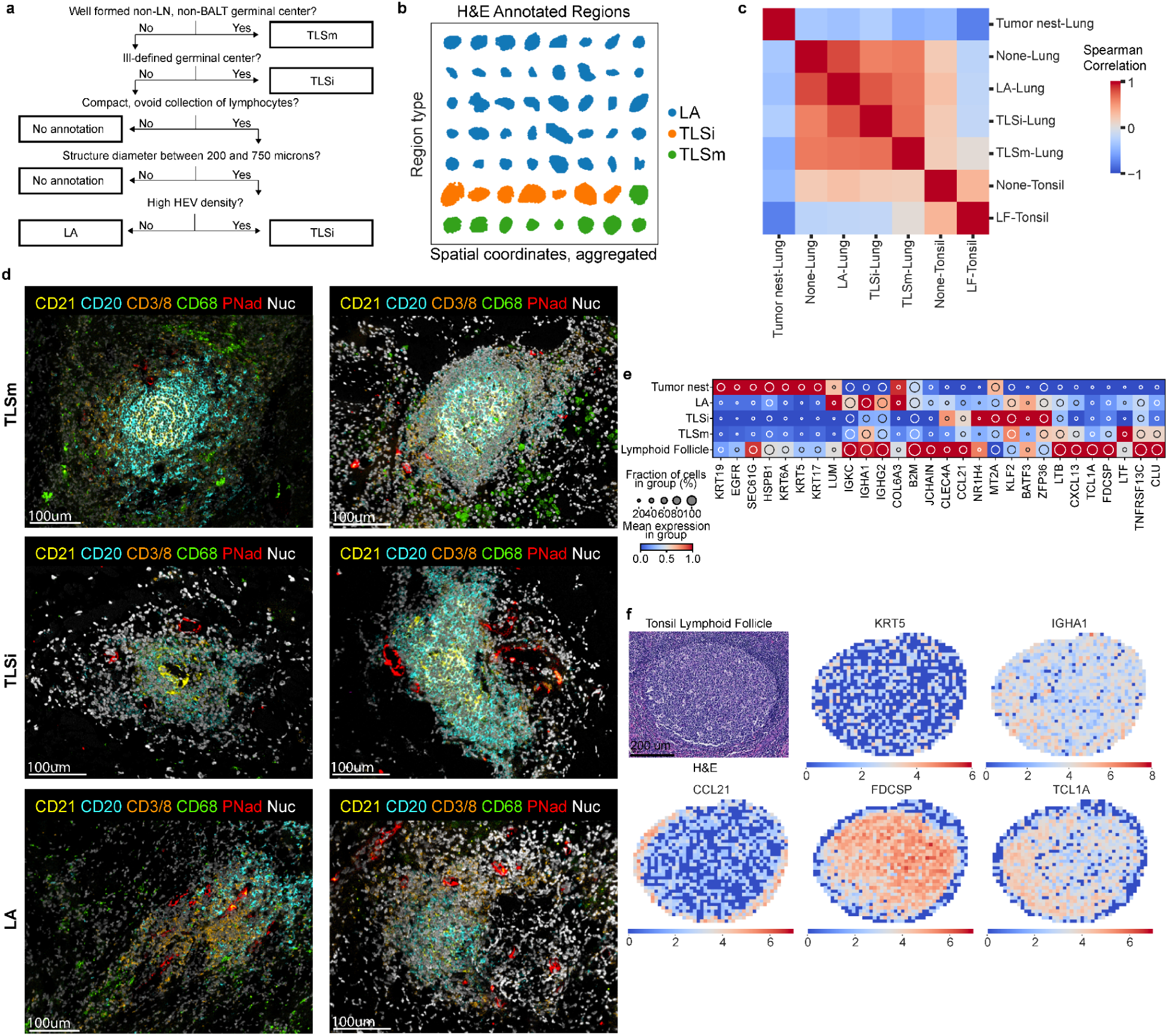
Overview of histopathological protocol, tonsil ROI, and MIBIscope spatial modality. **(A)** Flowchart for the histopathological annotation of ROI **(B)** Spatial plot of all initially annotated ROI demonstrating variance in size and morphology, annotated by classification into LA, TLSi, and TLSm **(C)** Spearman correlation of pseudobulked transcripts after log normalization across ROI histopathological classification and tissue source **(D)** MIBIscope spatial proteomics CD21 (FDCs), CD20 (B cells), CD3/8 (T cells), CD68 (Macrophages), PNad (HEVs, endothelial cells), and Nuc/DAPI (nuclei), scale bars = 100µm. **(E)** Dotplot depicting ROI-based differential gene expression by histological classification, values scaled after log normalization. Includes Tonsil Lymphoid Follicles. **(F)** Segmentation-free visualization of key differentially expressed genes *KRT5, IGHA1, CCL21, FDCSP*, and *TCL1A* within spatially-binned grids, with each FOV divided into 2,500 bins of about 258µm^2^ each. Transcripts were log normalized per spatial bin. For depicted tonsil lymphoid follicle, scale bar = 200µm.

**Extended Data Figure 2:**
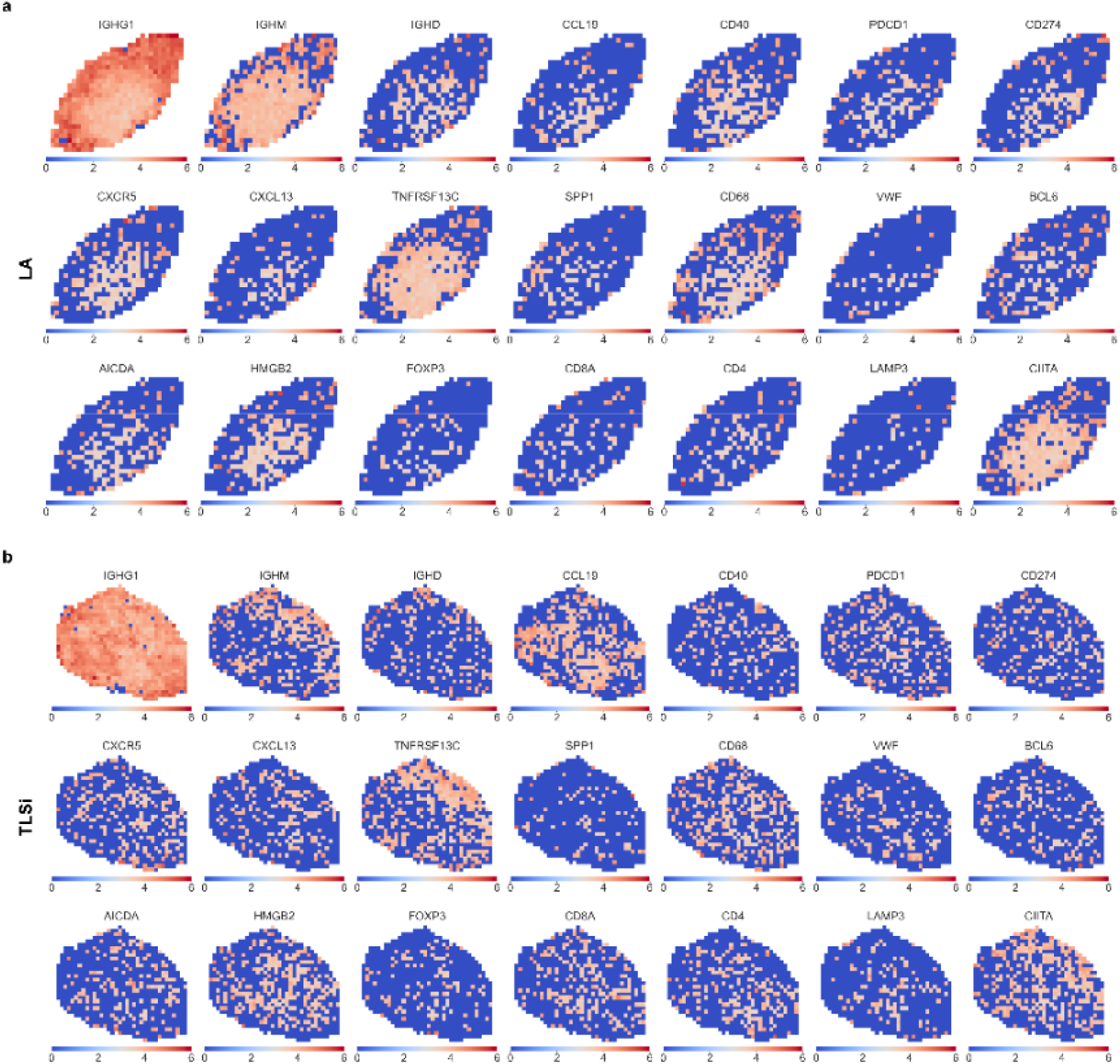
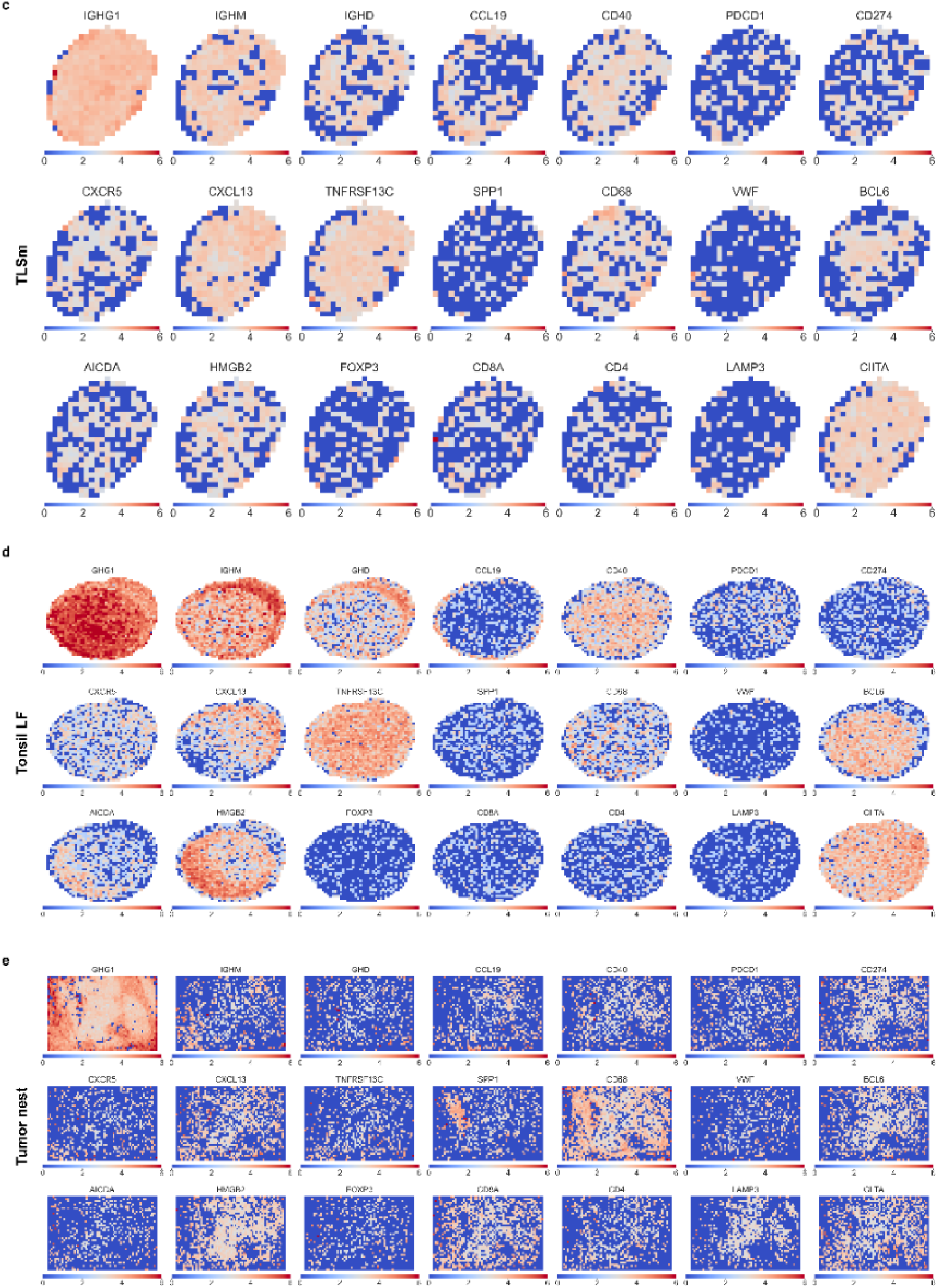
Segmentation-free visualization of key gene expression across representative ROI. Segmentation-free visualization of key marker genes (IGHG1, IGHM, IGHD, CCL19, CD40, PDCD1, CD274, CXCR5, CXCL13, TNFRSF13C, SPP1, CD68, VWF, BCL6, AICDA, HMGB2, FOXP3, CD8A, CD4, LAMP3, and CIITA) within spatially-binned grids, with each FOV divided into 2,500 bins of about 258µm^2^ each. **(A)** LA **(B)** TLSi **(C)** TLSm **(D)** Tonsil lymphoid follicle and **(E)** Tumor nests

**Extended Data Figure 3:**
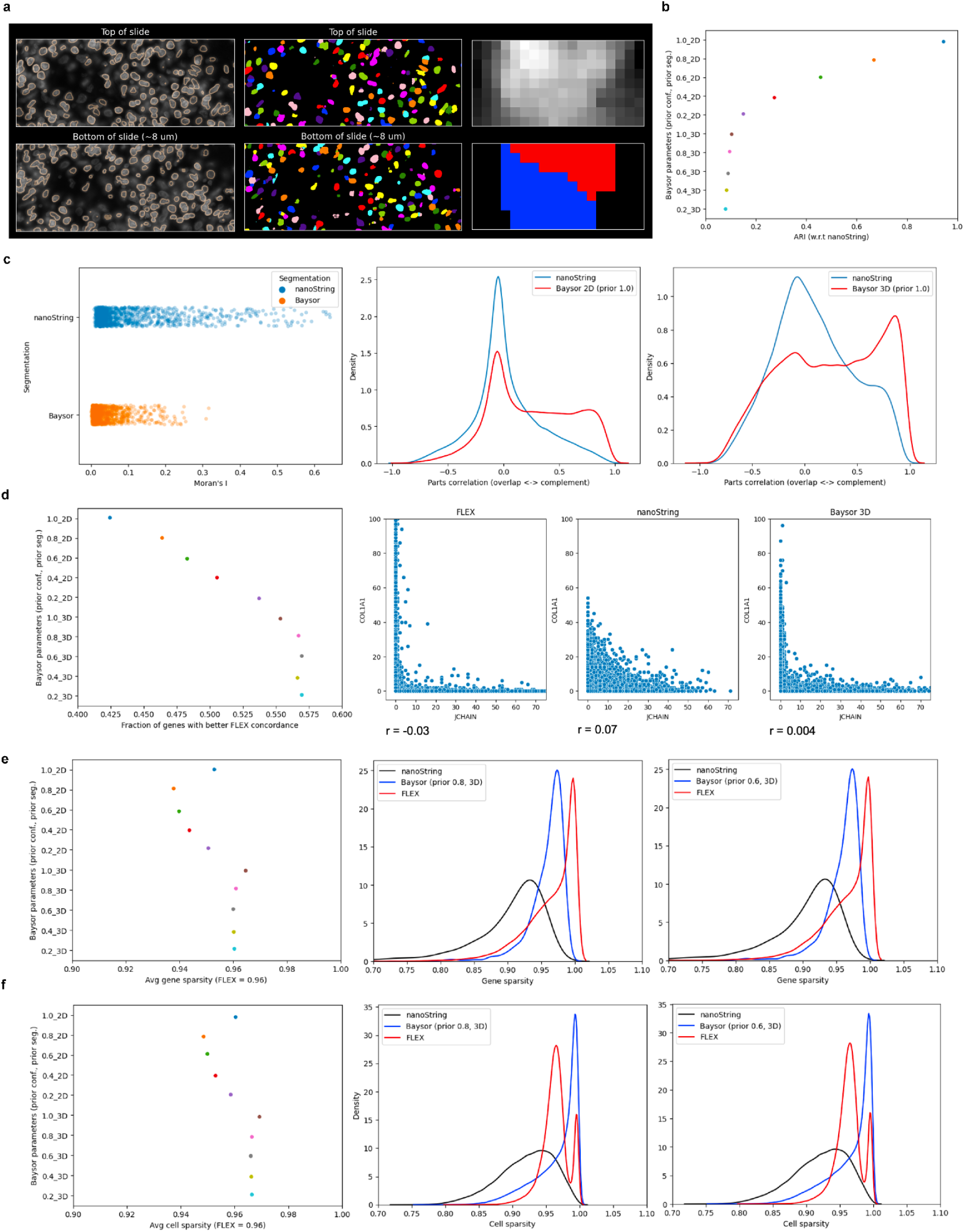
Evaluation of segmentation and transcript quantification using raw CosMx imaging data. **(A)** Visualization of z-stack images scanned by the CosMx SMI instrument showing the cell disparity between the top of the slide and bottom of the slide, approximately 8µm range. Segmentation mask outlines (left) with colors annotating unique cells (center) and a representative 3D stack of two distinct, horizontally coinciding cells denoted by color **(B)** Transcript ARI with respect to out-of-the-box cell/transcript assignments across 2D and 3D priors used for Baysor transcript assignment, showing a strong 2D prior closely resembles the out-of-the-box assignments **(C)** Strip plot of Moran’s I per gene, representing correlated expression between neighbors. High values interpreted as transcript spillover or mis-assignment (left). Density plots comparing the parts correlation value of the out-of-the-box transcript assignments to the 2D (center) and 3D (right) Baysor priors. Metric measures correlation between overlapping and complementary regions, where higher values indicate higher quality segmentation and transcript assignment. **(D)** Point plot showing CosMx-Flex gene correlation concordance metric calculated across different prior strengths and segmentation dimensions. Gene-gene correlations are calculated across cells in CosMx and Flex independently, ranked, and compared via Spearman’s rho. Measures the similarity in pairwise gene-gene correlations across cells. First, pairwise correlation coefficients (CC) are calculated for each gene-gene pair across cells (independently for both CosMX and Flex). Segmentation errors will reduce this correlation relative to Flex (left). Scatterplots comparing two genes which should not be coexpressed, COL1A1 (mesenchymal) and JCHAIN (plasma cell), across representative modalities and transcript-assignment methods (right). **(E)** Point plot showing average Gini coefficient for each gene across all cells (left); with 0 interpreted as gene expression evenly distributed across all cells and 1 interpreted as genes being only expressed in one cell. Full coefficient distribution for 0.8 (center) and 0.6 (right) 3D prior strengths shown. A low cell sparsity in CosMX compared to Flex indicates gene spillover and segmentation errors. **(F)** Point plot showing average Gini coefficient for each gene within the same cells (left); a value of 0 is interpreted as all cells expressing all genes in an evenly distributed manner. Contrasting this, 1 is observed for cells that express only one gene. Full coefficient distribution for 0.8 (center) and 0.6 (right) 3D prior strengths shown.

**Extended Data Figure 4:**
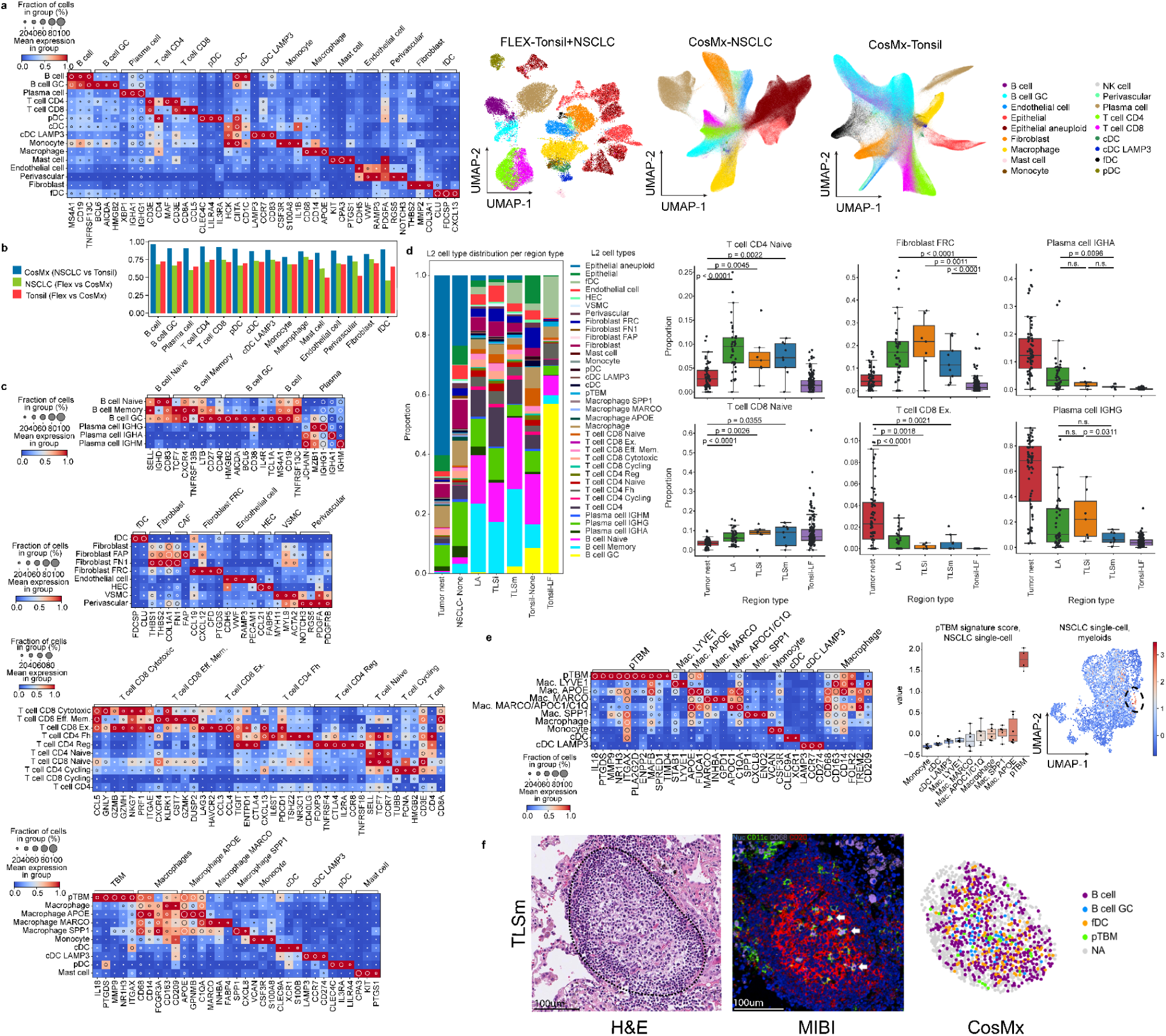
Comprehensive L1 and L2 cell subtype gene expression characterization and detection within ROI. **(A)** Tonsil CosMx (left) UMAPs (right) **(B)** Pseudobulked spearman correlation per cell type **(C)** NSCLC CosMx Cell type-specific L2 marker gene expression. values scaled after log normalization **(D)** Stacked barplot showing proportional representation of L2 cell types captured across all classified ROI (left). Boxplots depicting L2 cell subtype representation, with proportions normalized by cell lineage per ROI. Mann-Whitney U tests were performed between ROI classifications per subtype (right). **(E)** Dotplot depicting high-resolution myeloid cell subtypes within NSCLC-derived SN and ST datasets, highlighting pTBM marker expression in NSCLC Flex dataset; values scaled after log normalization (left). Mean pTBM signature score per myeloid subtype per sample, calculated on log normalized values (center) UMAP depicting NSCLC Flex myeloid cells, pTBM signature score overlaid (right) **(F)** Representative TLSm depicted across three serially-sliced modalities. H&E, scale bar = 100µm (left). MIBIscope with Nuc/DAPI, nuclei; CD11c, pTBM/Macrophages; CD68, Macrophages; CD20, B cells (center). CosMx, overlaid with L2 cell subtype annotations (right), scale bar = 100µm.

**Extended Data Figure 5:**
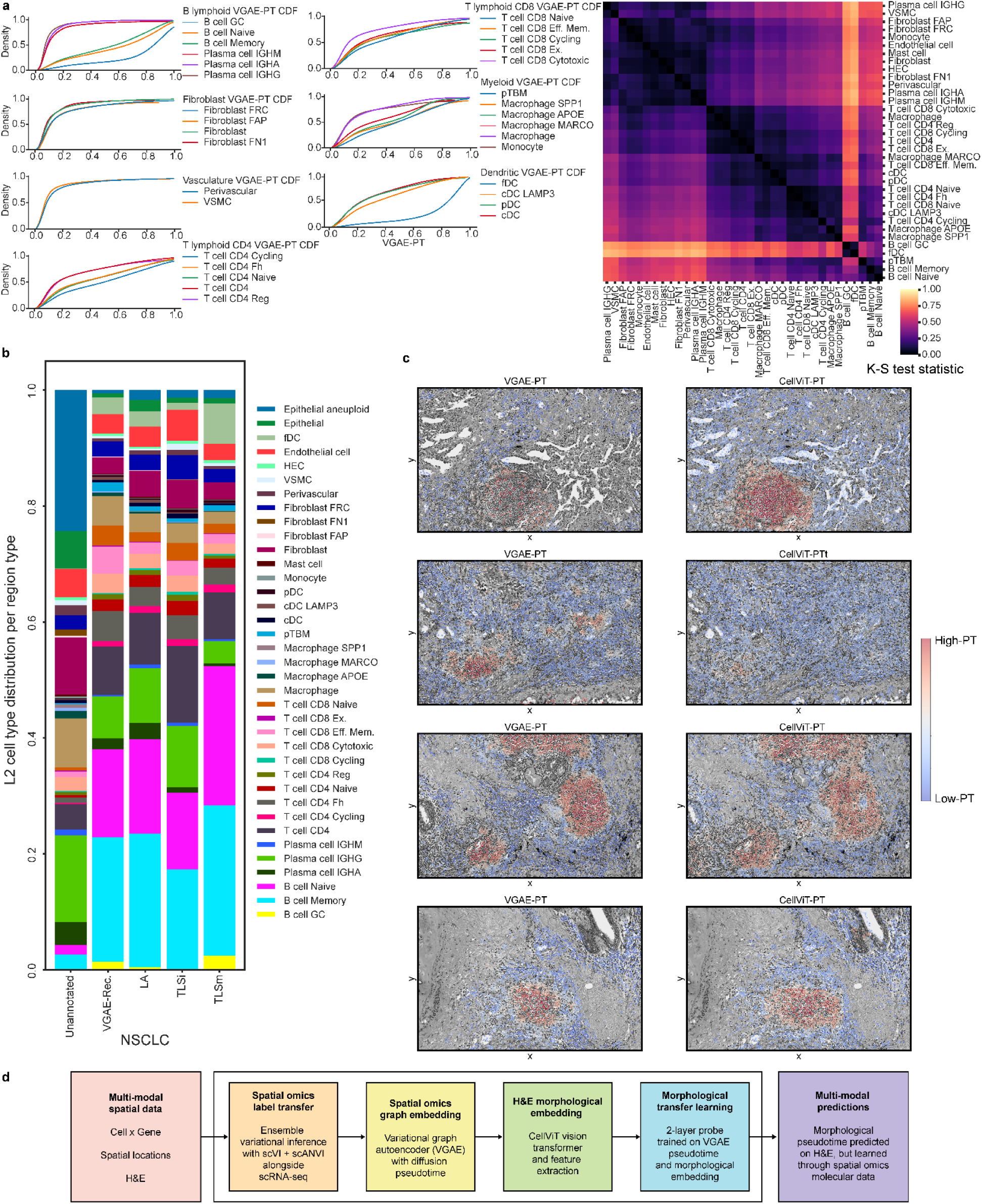
VGAE-PT distributions across all L2 cell subtypes, VGAE-Rec. ROI, and relationship with CellViT-PT. **(A)** CDF of VGAE-PT values partitioned by L2 cell subtype (left). Corresponding K-S test statistic shown for each curve, compared pairwise, with full matrix in Supplementary Table 7 (right). **(B)** Stacked barplot showing proportional representation of L2 cell types captured across all classified ROI, in addition to VGAE-Rec. ROI. **(C)** Representative spatial overlays of VGAE-PT compared to CellViT-PT **(D)** Schematic showing step-wise data input, processing steps, and output for cell type annotation, graph embedding, H&E morphological feature embedding, and performing multi-modal predictions to generate pseudotime based on the H&E morphological features.

**Extended Data Figure 6:**
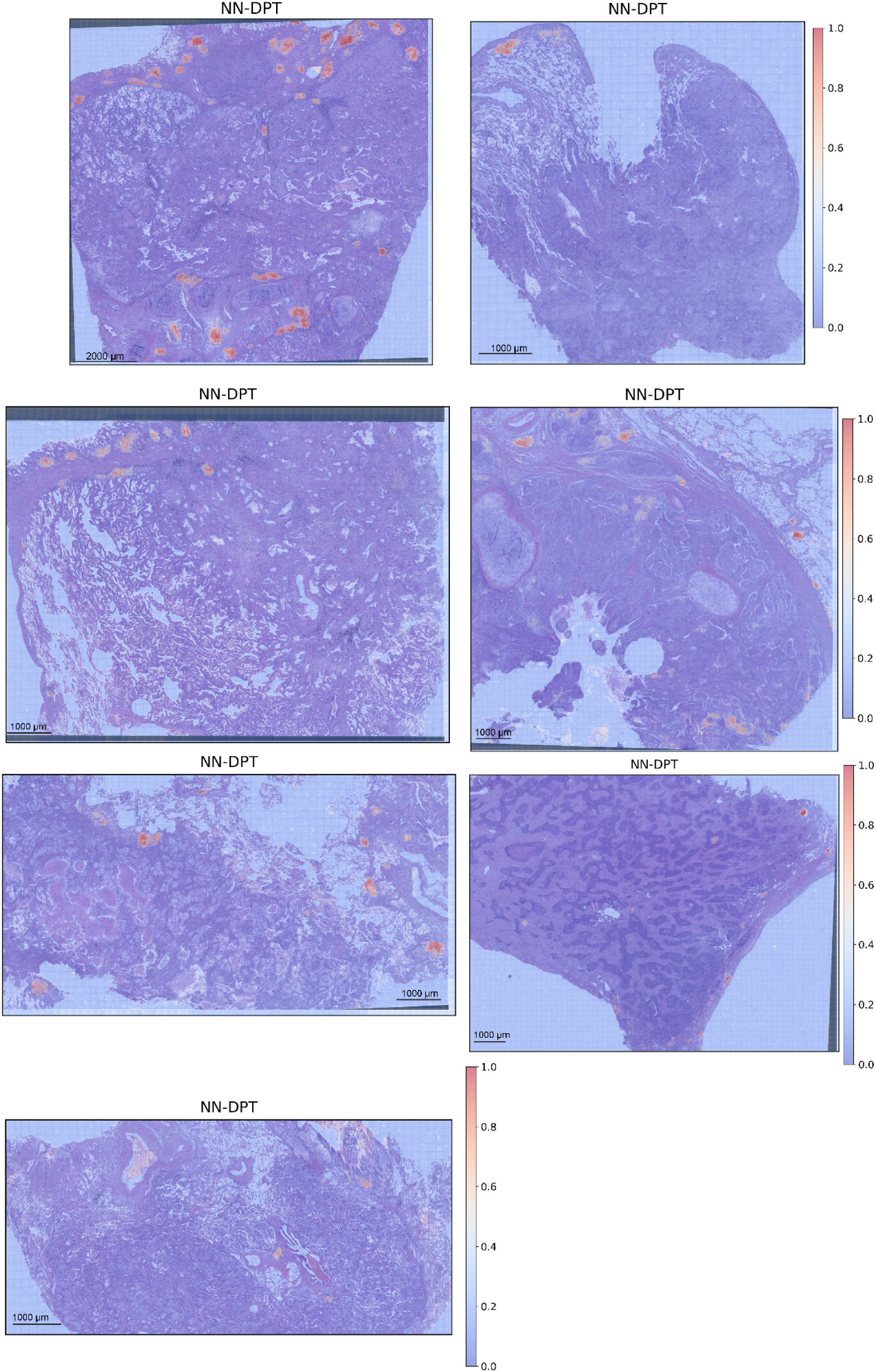
Whole slide overlays of predicted CellViT-PT. Predicted CellViT morphological pseudotime values across whole H&E slide images after slide alignment to CosMx FOVs and training on spatially registered VGAE-PT values. Scale bar varies between slides, between 1000 to 2000 µm.

**Extended Data Figure 7:**
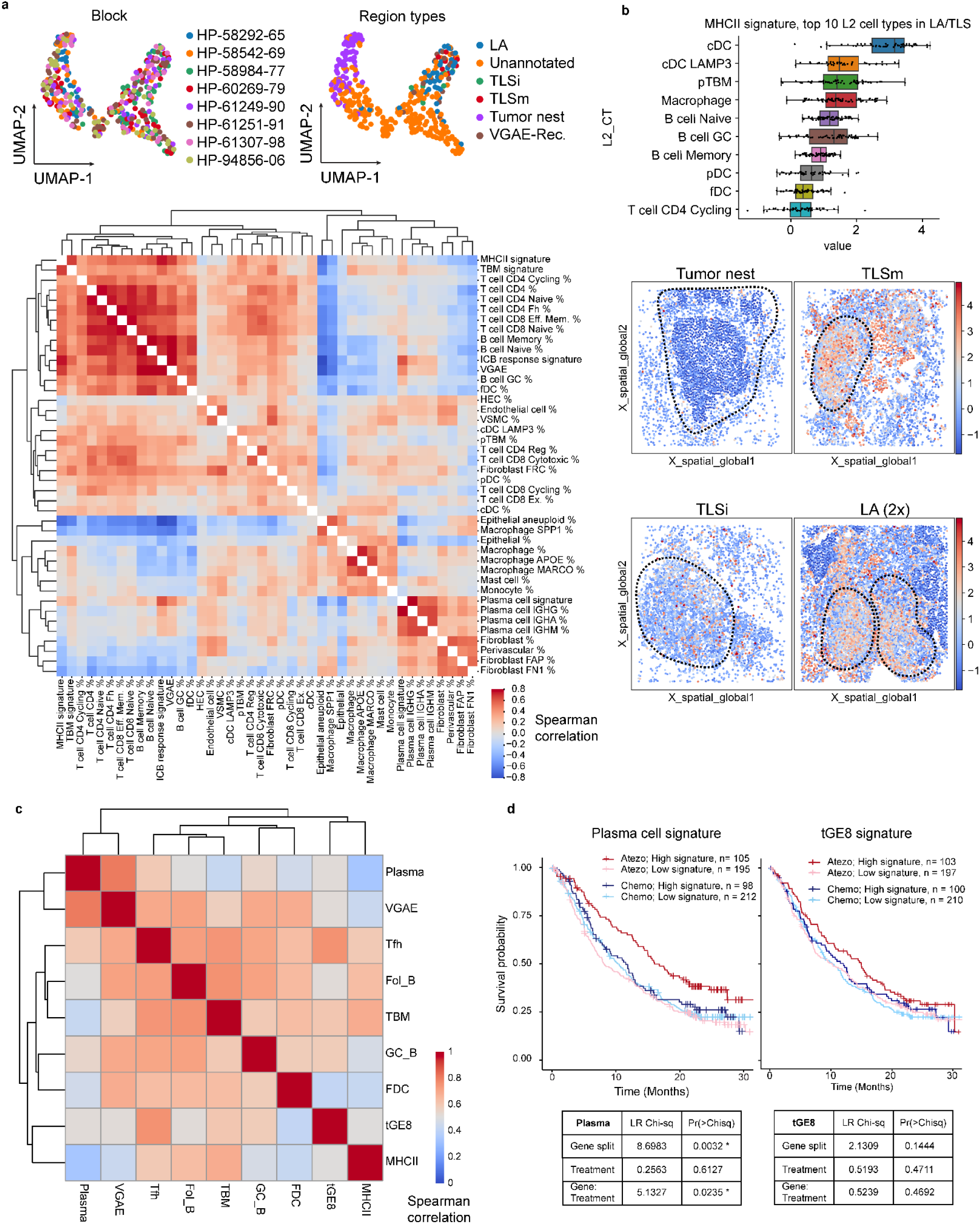
Spatial relationships of gene signature scores and contribution to patient survival in OAK. **(A)** UMAP of ROI by L2 cell subtype composition demonstrating gene signature score representation within ROI alongside cell subtype proportions (upper). Spearman correlation matrix of ROI mean signature scores and proportional representations of L2 cell subtypes (lower) **(B)** MHCII signature score averaged per L2 cell subtype and ROI, only including LA, TLSi, and TLSm (upper). Spatial plots depicting localized enrichments of MHCII signature scores across classified ROI (lower) **(C)** Spearman correlations calculated in OAK, of gene signatures applied to survival analyses across OAK and TCGA **(D)** Kaplan-Meier curves for plasma cell and tGE8 signature in OAK. Tables contain ANOVA results with 1 degree of freedom, full results in Supplementary Table 10.

**Extended Data Figure 8:**
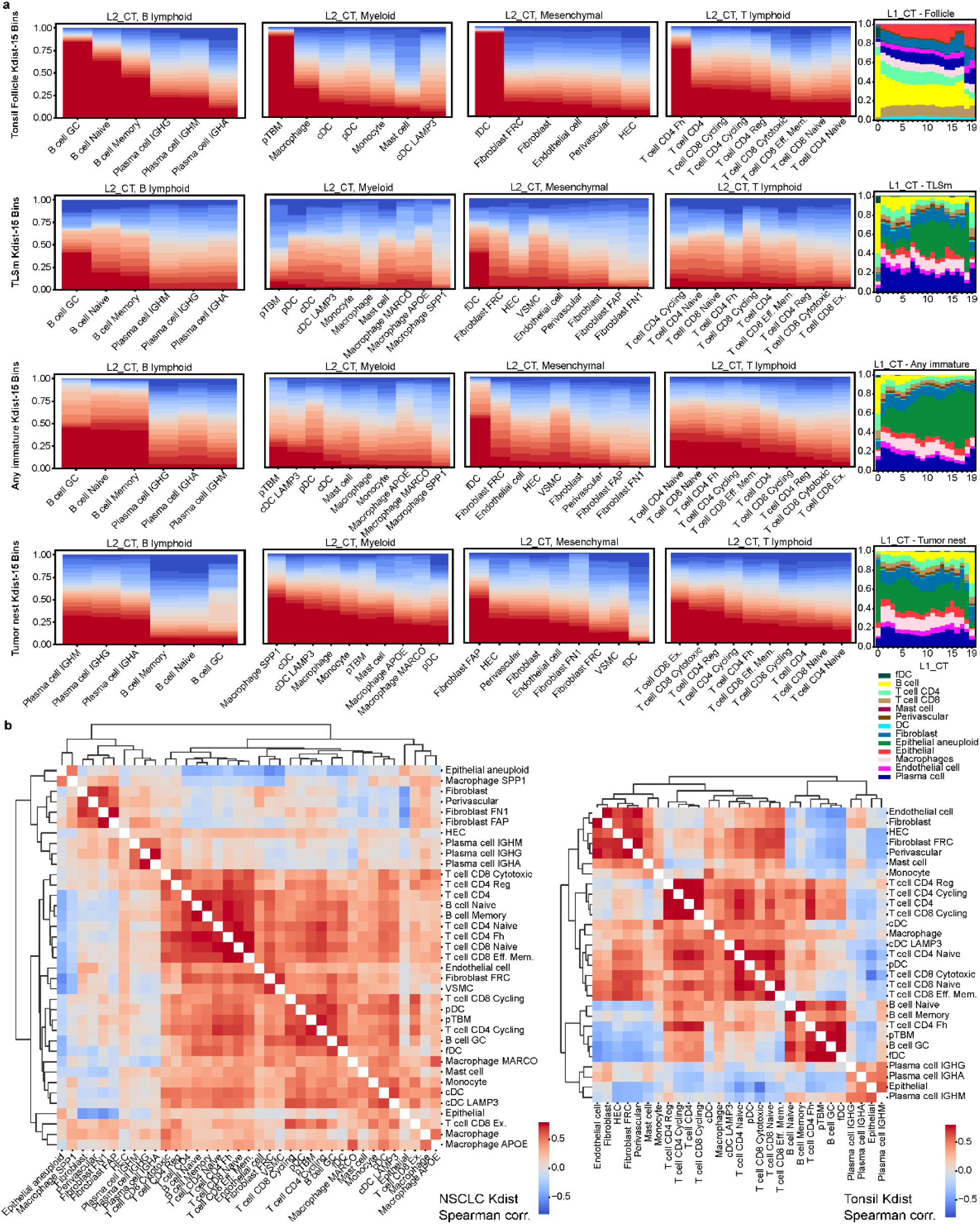
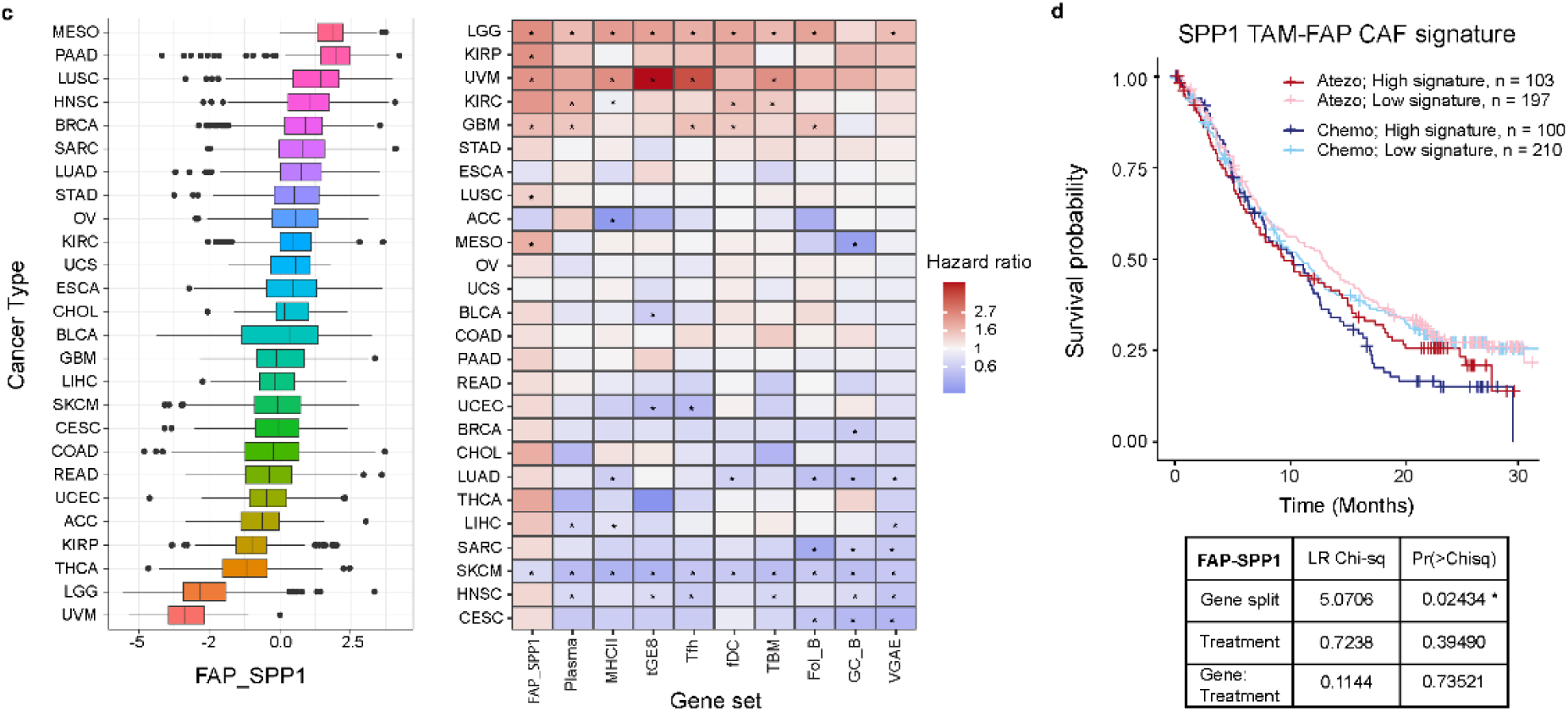
Spatial distance analyses partitioned by cell lineage and ROI classification alongside corresponding survival analysis. **(A)** Stacked barplots visualizing L2 cell subtype representation across 20 K-dist bins calculated on the basis of each individual tissue block, barplots grouped by cell lineage. Cells within the boundaries of ROI were placed in the nearest bin, shown in red (left). Bin-wise stacked barplots showing proportional representation of L1 cell types (right). **(B)** Full K-dist co-localization matrix. Spearman correlation values were calculated across all tissue blocks, hierarchically clustered, and grouped by tissue source. **(C)** Boxplot of GSVA scores calculated for FAP-SPP1 gene signature across TCGA cancer types (left). Overall survival and hazard ratios given GSVA gene set scores quantizing patient groups, asterisks indicate p < 0.05 (right) **(D)** Kaplan-Meier plot for the FAP-SPP1 gene signatures scored by GSVA in the OAK cohort, dichotomization procedure detailed in methods. Tables contain ANOVA results with 1 degree of freedom, full results in Supplementary Table 10.

