## Extended_data_figures for "Spatial Decoding of Tertiary Lymphoid Structure Maturation in Non-Small Cell Lung Cancer Using Deep Neural Networks"

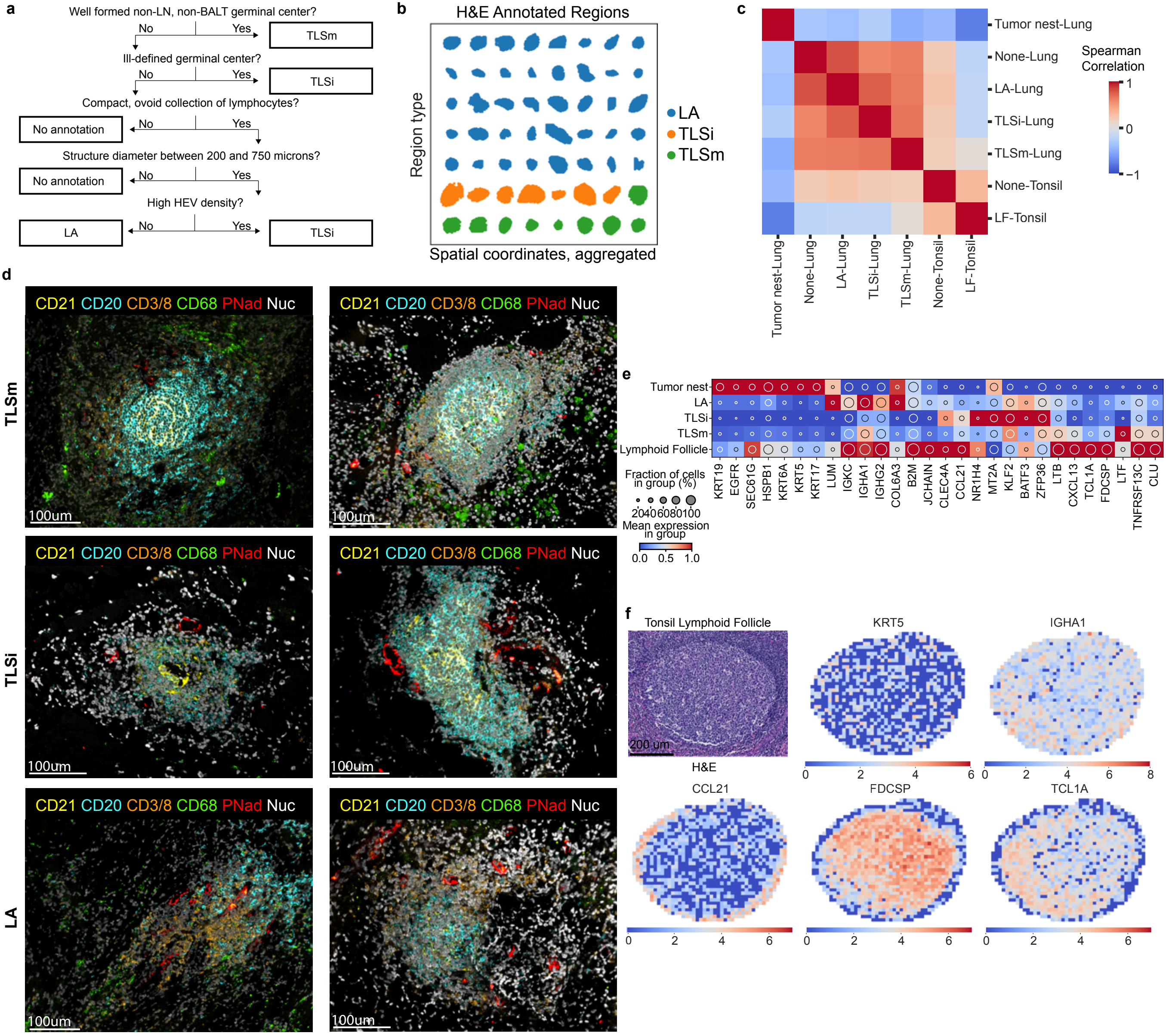
a
Well formed non-LN, non-BALT germinal center?
No
Yes
TLSm
Ill-defined germinal center?
No
Yes
TLSi
Compact, ovoid collection of lymphocytes?
No annotation
No
Yes
Structure diameter between 200 and 750 microns?
No annotation
No
Yes
High HEV density?
LA
No
Yes
TLSi
b
H&E Annotated Regions
Region type
Spatial coordinates, aggregated
LA
TLSi
TLSm
c
Tumor nest-Lung
None-Lung
LA-Lung
TLSi-Lung
TLSm-Lung
None-Tonsil
LF-Tonsil
Spearman Correlation
1
0
−1
d
CD21 CD20 CD3/8 CD68 PNad Nuc
TLSm
TLSi
LA
100um
e
Tumor nest
LA
TLSi
TLSm
Lymphoid Follicle
KRT19 EGFR SEC61G HSPB1 KRT6A KRT5 KRT17 LUM IGKC IGHA1 IGHG2 COL6A3 B2M JCHAIN CLEC4A CCL21 NR1H4 MT2A KLF2 BATF3 ZFP36 LTB CXCL13 TCL1A FDCSP LTF TNFRSF13C CLU
Fraction of cells in group (%)
20 40 60 80 100
Mean expression in group
0.0 0.5 1.0
f
Tonsil Lymphoid Follicle
200 um
H&E
KRT5
IGHA1
CCL21
FDCSP
TCL1A
0 2 4 6
0 2 4 6 8

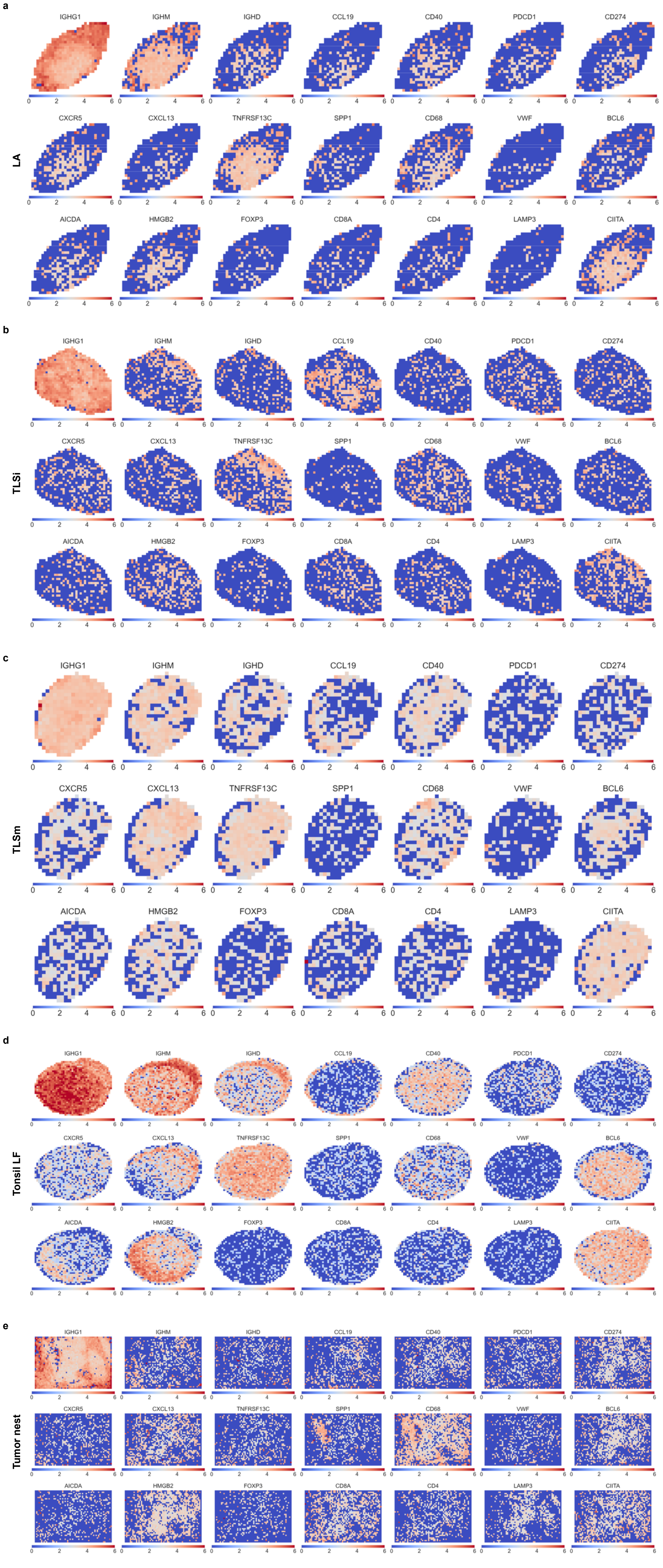
a
LA
b
TLSi
c
TLSm
d
Tonsil LF
e
Tumor nest
IGHG1
IGHM
IGHD
CCL19
CD40
PDCD1
CD274
CXCR5
CXCL13
TNFRSF13C
SPP1
CD68
VWF
BCL6
AICDA
HMGB2
FOXP3
CD8A
CD4
LAMP3
CIITA
0
2
4
6

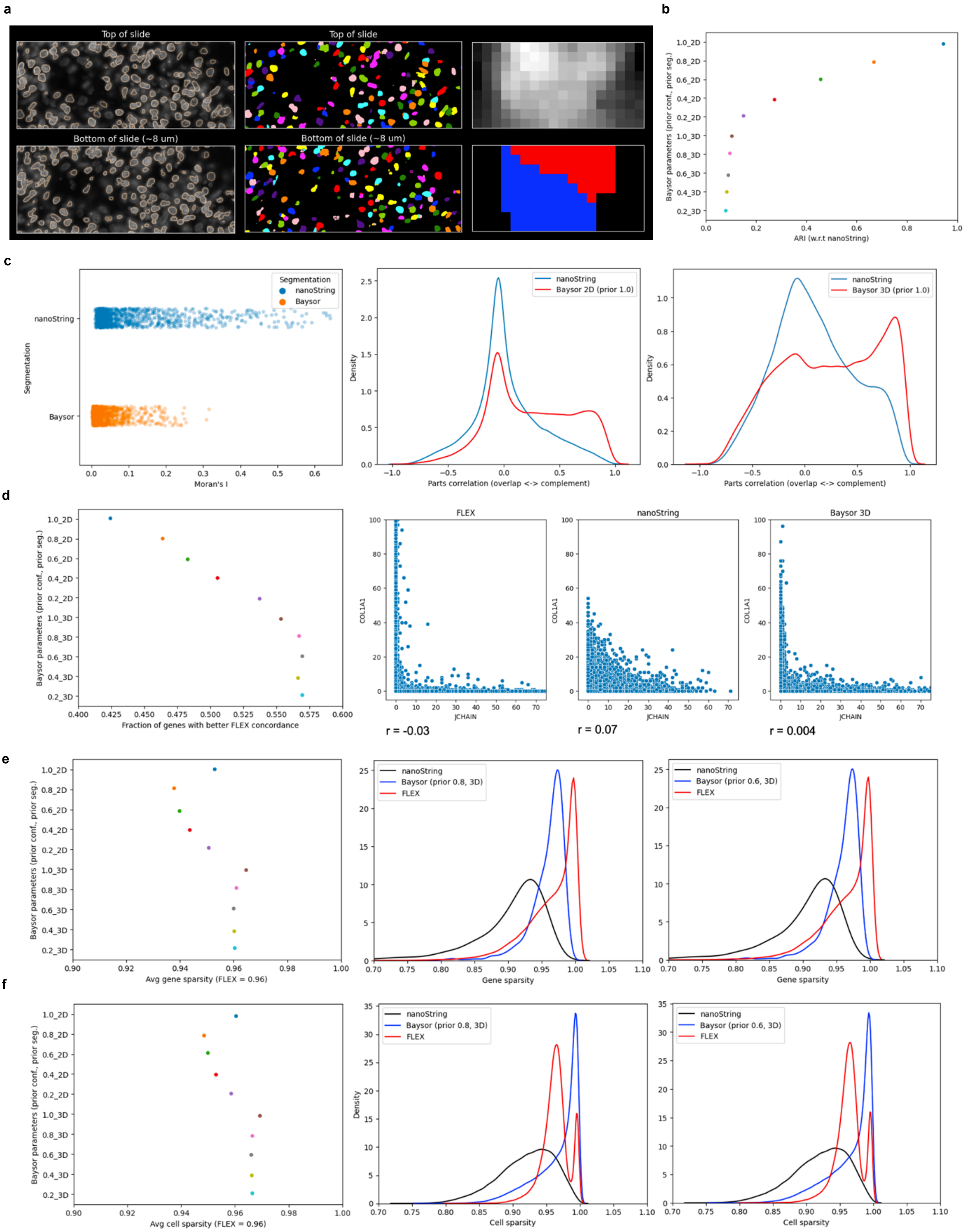

a
Top of slide
Top of slide
Bottom of slide (~8 um)
Bottom of slide (~8 um)
b
Baysor parameters (prior conf., prior seg.)
1.0_2D
0.8_2D
0.6_2D
0.4_2D
0.2_2D
1.0_3D
0.8_3D
0.6_3D
0.4_3D
0.2_3D
ARI (w.r.t nanoString)
c
Segmentation
nanoString
Baysor
Moran's I
Density
nanoString
Baysor 2D (prior 1.0)
Parts correlation (overlap <-> complement)
Baysor 3D (prior 1.0)
d
Fraction of genes with better FLEX concordance
FLEX
nanoString
Baysor 3D
COL1A1
JCHAIN
r = -0.03
r = 0.07
r = 0.004
e
Avg gene sparsity (FLEX = 0.96)
Baysor (prior 0.8, 3D)
Baysor (prior 0.6, 3D)
FLEX
Gene sparsity
f
Avg cell sparsity (FLEX = 0.96)
Cell sparsity

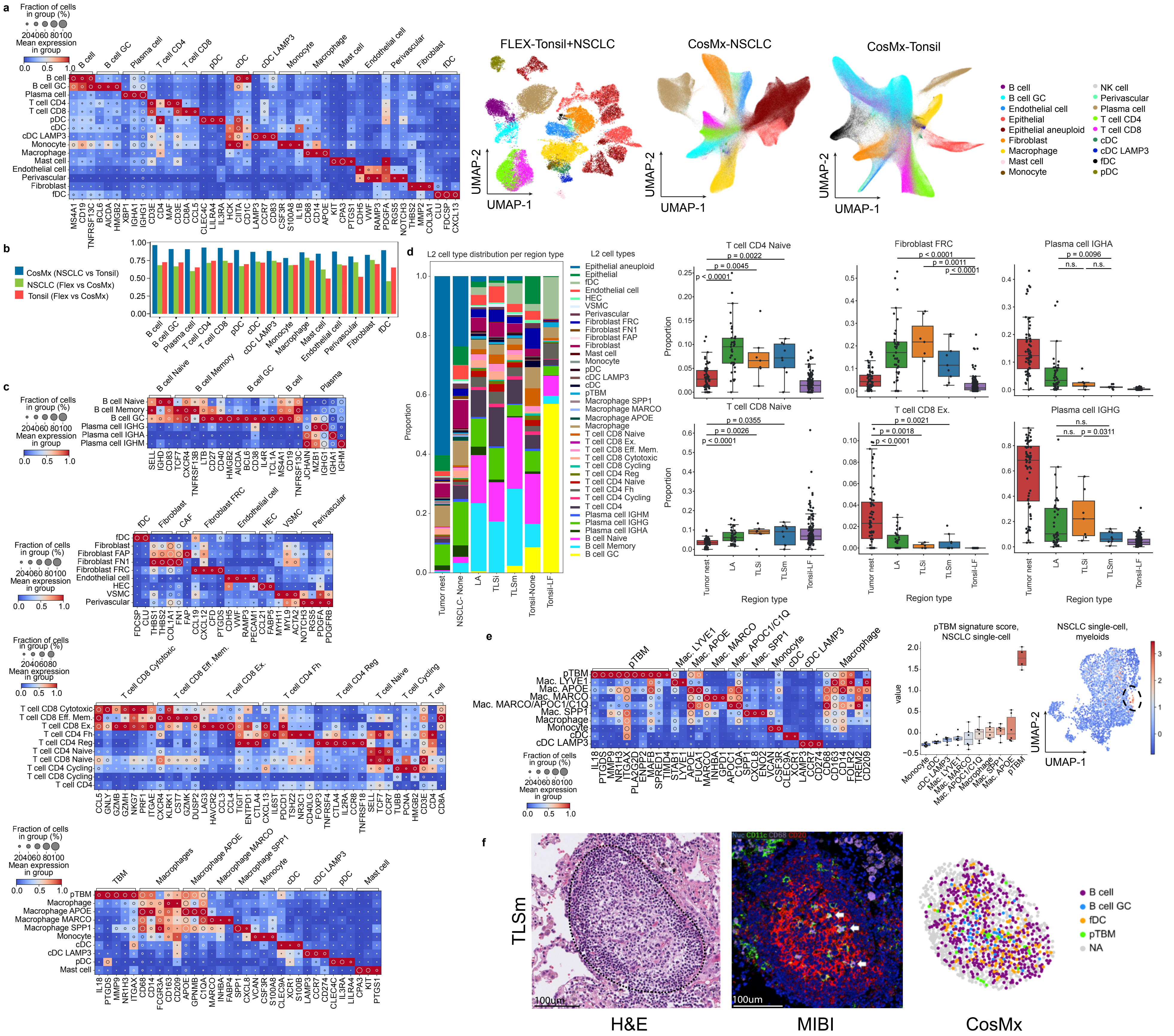
a
Fraction of cells in group (%)
20 40 60 80 100
Mean expression in group
0.0 0.5 1.0
B cell
B cell GC
Plasma cell
T cell CD4
T cell CD8
pDC
cDC
cDC LAMP3
Monocyte
Macrophage
Mast cell
Endothelial cell
Perivascular
Fibroblast
fDC
MS4A1 CD19 TNFRSF13C BCL6 AICDA HMGB2 XBP1 IGHA1 IGHG1 CD3E CD4 MAF CD3E CD8A CCL5 CLEC4C LILRA4 IL3RA HCK CIITA CD1C LAMP3 CCR7 CD83 CSF3R S100A8 IL1B CD68 CD14 APOE KIT CPA3 PTGS1 CDH5 VWF RAMP3 PDGFA RGS5 NOTCH3 THBS2 MMP2 COL3A1 CLU FDCSP CXCL13
FLEX-Tonsil+NSCLC
CosMx-NSCLC
CosMx-Tonsil
UMAP-1
UMAP-2
B cell
B cell GC
Endothelial cell
Epithelial
Epithelial aneuploid
Fibroblast
Macrophage
Mast cell
Monocyte
NK cell
Perivascular
Plasma cell
T cell CD4
T cell CD8
cDC
cDC LAMP3
fDC
pDC
b
CosMx (NSCLC vs Tonsil)
NSCLC (Flex vs CosMx)
Tonsil (Flex vs CosMx)
1.00
0.75
0.50
0.25
0.00
B cell
B cell GC
Plasma cell
T cell CD4
T cell CD8
pDC
cDC
cDC LAMP3
Monocyte
Macrophage
Mast cell
Endothelial cell
Perivascular
Fibroblast
fDC
c
B cell Naive
B cell Memory
B cell GC
B cell
Plasma
Plasma cell IGHG
Plasma cell IGHA
Plasma cell IGHM
SELL IGHD CD83 TCF7 CXCR4 TNFRSF13B LTB CD27 CD40 HMGB2 AICDA BCL6 CD38 IL4R TCL1A MS4A1 CD19 TNFRSF13C JCHAIN MZB1 IGHG1 IGHA1 IGHM
fDC
Fibroblast
CAF
Fibroblast FRC
Endothelial cell
HEC
VSMC
Perivascular
Fibroblast FAP
Fibroblast FN1
FDCSP CLU THBS1 THBS2 COL1A1 FN1 FAP CCL19 CXCL12 CFD PTGDS CDH5 VWF RAMP3 PECAM1 CCL21 FABP5 MYH11 MYL9 ACTA2 NOTCH3 RGS5 PDGFA PDGFRB
T cell CD8 Cytotoxic
T cell CD8 Eff. Mem.
T cell CD8 Ex.
T cell CD4 Fh
T cell CD4 Reg
T cell Naive
T cell Cycling
T cell
T cell CD4 Naive
T cell CD8 Naive
T cell CD4 Cycling
T cell CD8 Cycling
T cell CD4
CCL5 GNLY GZMB GZMH NKG7 PRF1 ITGAE CXCR4 KLRK1 CST7 GZMK DUSP2 LAG3 HAVCR2 CCL3 CCL4 TIGIT ENTPD1 CTLA4 CXCL13 IL6ST PDCD1 TSHZ2 NR3C1 CD40LG FOXP3 TNFRSF4 CTLA4 IL2RA CCR8 TNFRSF18 SELL TCF7 CCR7 TUBB PCNA HMGB2 CD3E CD4 CD8A
pTBM
TBM
Macrophages
Macrophage
Macrophage APOE
Macrophage MARCO
Macrophage SPP1
Monocyte
cDC
cDC LAMP3
pDC
Mast cell
IL18 PTGDS MMP9 NR1H3 ITGAX CD68 CD14 FCGR3A CD163 CD209 APOE GPNMB C1QA MARCO INHBA FABP4 SPP1 CXCL8 VCAN CSF3R S100A8 CLEC9A XCR1 S100B LAMP3 CCR7 CD274 CLEC4C IL3RA LILRA4 CPA3 KIT PTGS1
d
L2 cell type distribution per region type
Proportion
Tumor nest
NSCLC- None
LA
TLSi
TLSm
Tonsil-None
Tonsil-LF
L2 cell types
Epithelial aneuploid
Epithelial
fDC
Endothelial cell
HEC
VSMC
Perivascular
Fibroblast FRC
Fibroblast FN1
Fibroblast FAP
Fibroblast
Mast cell
Monocyte
pDC
cDC LAMP3
cDC
pTBM
Macrophage SPP1
Macrophage MARCO
Macrophage APOE
Macrophage
T cell CD8 Naive
T cell CD8 Ex.
T cell CD8 Eff. Mem.
T cell CD8 Cytotoxic
T cell CD8 Cycling
T cell CD4 Reg
T cell CD4 Naive
T cell CD4 Fh
T cell CD4 Cycling
T cell CD4
Plasma cell IGHM
Plasma cell IGHG
Plasma cell IGHA
B cell Naive
B cell Memory
B cell GC
T cell CD4 Naive
p = 0.0022
p = 0.0045
p < 0.0001
Fibroblast FRC
p < 0.0001
p = 0.0011
p < 0.0001
Plasma cell IGHA
p = 0.0096
n.s.
n.s.
T cell CD8 Naive
p = 0.0355
p = 0.0026
p < 0.0001
T cell CD8 Ex.
p = 0.0021
p = 0.0018
p < 0.0001
Plasma cell IGHG
n.s.
n.s.
p = 0.0311
Region type
e
pTBM
Mac. LYVE1
Mac. APOE
Mac. MARCO
Mac. APOC1/C1Q
Mac. SPP1
Monocyte
cDC
cDC LAMP3
Macrophage
Mac. MARCO/APOC1/C1Q
IL18 PTGDS MMP9 NR1H3 ITGAX PLA2G2D ENPP2 MAFB SPRED1 TIMD4 STAB1 LYVE1 APOE FUCA1 MARCO INHBA GPD1 APOC1 C1QA SPP1 CXCL8 ENO2 VCAN CSF3R CLEC9A XCR1 LAMP3 CCR7 CD274 CD68 CD163 CD14 FOLR2 TREM2 CD209
pTBM signature score, NSCLC single-cell
value
NSCLC single-cell, myeloids
UMAP-1
UMAP-2
f
TLSm
Nuc CD11c CD68 CD20
100um
H&E
MIBI
CosMx
B cell
B cell GC
fDC
pTBM
NA

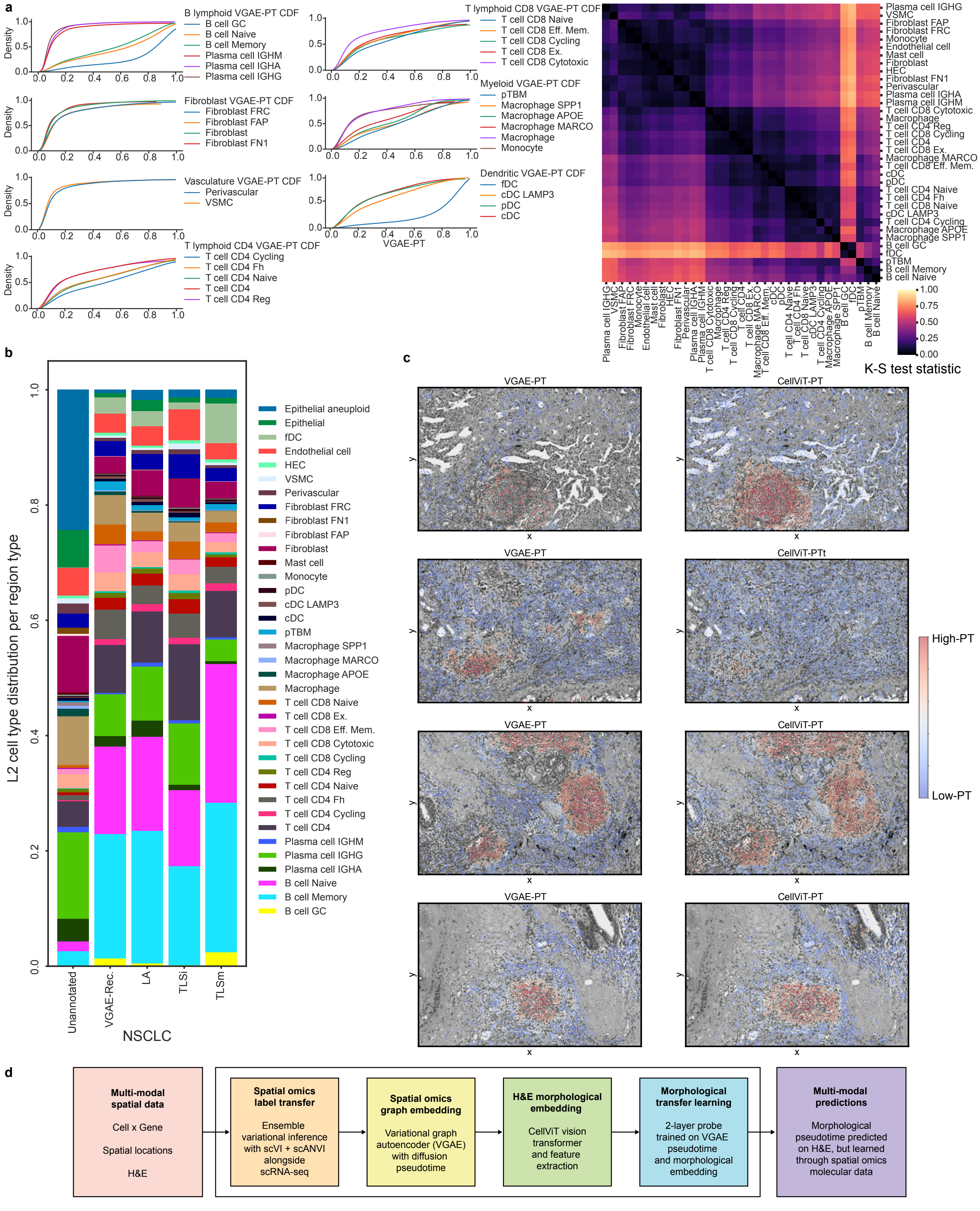
a
B lymphoid VGAE-PT CDF
B cell GC
B cell Naive
B cell Memory
Plasma cell IGHM
Plasma cell IGHA
Plasma cell IGHG
Fibroblast VGAE-PT CDF
Fibroblast FRC
Fibroblast FAP
Fibroblast
Fibroblast FN1
Vasculature VGAE-PT CDF
Perivascular
VSMC
T lymphoid CD4 VGAE-PT CDF
T cell CD4 Cycling
T cell CD4 Fh
T cell CD4 Naive
T cell CD4
T cell CD4 Reg
T lymphoid CD8 VGAE-PT CDF
T cell CD8 Naive
T cell CD8 Eff. Mem.
T cell CD8 Cycling
T cell CD8 Ex.
T cell CD8 Cytotoxic
Myeloid VGAE-PT CDF
pTBM
Macrophage SPP1
Macrophage APOE
Macrophage MARCO
Macrophage
Monocyte
Dendritic VGAE-PT CDF
fDC
cDC LAMP3
pDC
cDC
Density
VGAE-PT
K-S test statistic
b
L2 cell type distribution per region type
Unannotated
VGAE-Rec.
LA
TLSi
TLSm
NSCLC
Epithelial aneuploid
Epithelial
fDC
Endothelial cell
HEC
VSMC
Perivascular
Fibroblast FRC
Fibroblast FN1
Fibroblast FAP
Fibroblast
Mast cell
Monocyte
pDC
cDC LAMP3
cDC
pTBM
Macrophage SPP1
Macrophage MARCO
Macrophage APOE
Macrophage
T cell CD8 Naive
T cell CD8 Ex.
T cell CD8 Eff. Mem.
T cell CD8 Cytotoxic
T cell CD8 Cycling
T cell CD4 Reg
T cell CD4 Naive
T cell CD4 Fh
T cell CD4 Cycling
T cell CD4
Plasma cell IGHM
Plasma cell IGHG
Plasma cell IGHA
B cell Naive
B cell Memory
B cell GC
c
VGAE-PT
CellViT-PT
CellViT-PTt
High-PT
Low-PT
d
Multi-modal spatial data
Cell x Gene
Spatial locations
H&E
Spatial omics label transfer
Ensemble variational inference with scVI + scANVI alongside scRNA-seq
Spatial omics graph embedding
Variational graph autoencoder (VGAE) with diffusion pseudotime
H&E morphological embedding
CellViT vision transformer and feature extraction
Morphological transfer learning
2-layer probe trained on VGAE pseudotime and morphological embedding
Multi-modal predictions
Morphological pseudotime predicted on H&E, but learned through spatial omics molecular data

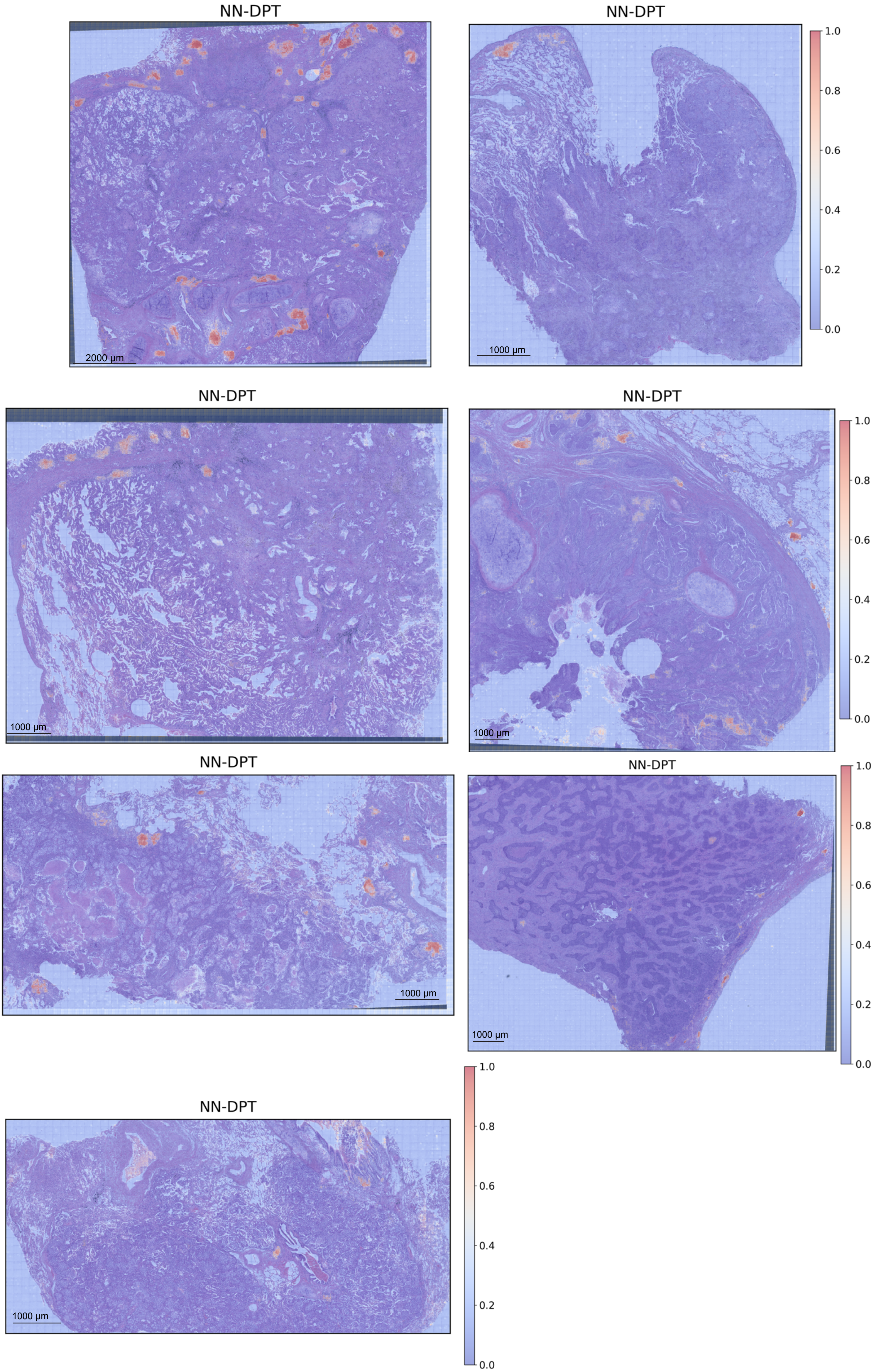
NN-DPT
2000 µm
NN-DPT
1000 µm
1.0
0.8
0.6
0.4
0.2
0.0
NN-DPT
1000 µm
NN-DPT
1000 µm
1.0
0.8
0.6
0.4
0.2
0.0
NN-DPT
1000 µm
NN-DPT
1000 µm
1.0
0.8
0.6
0.4
0.2
0.0
NN-DPT
1000 µm
1.0
0.8
0.6
0.4
0.2
0.0

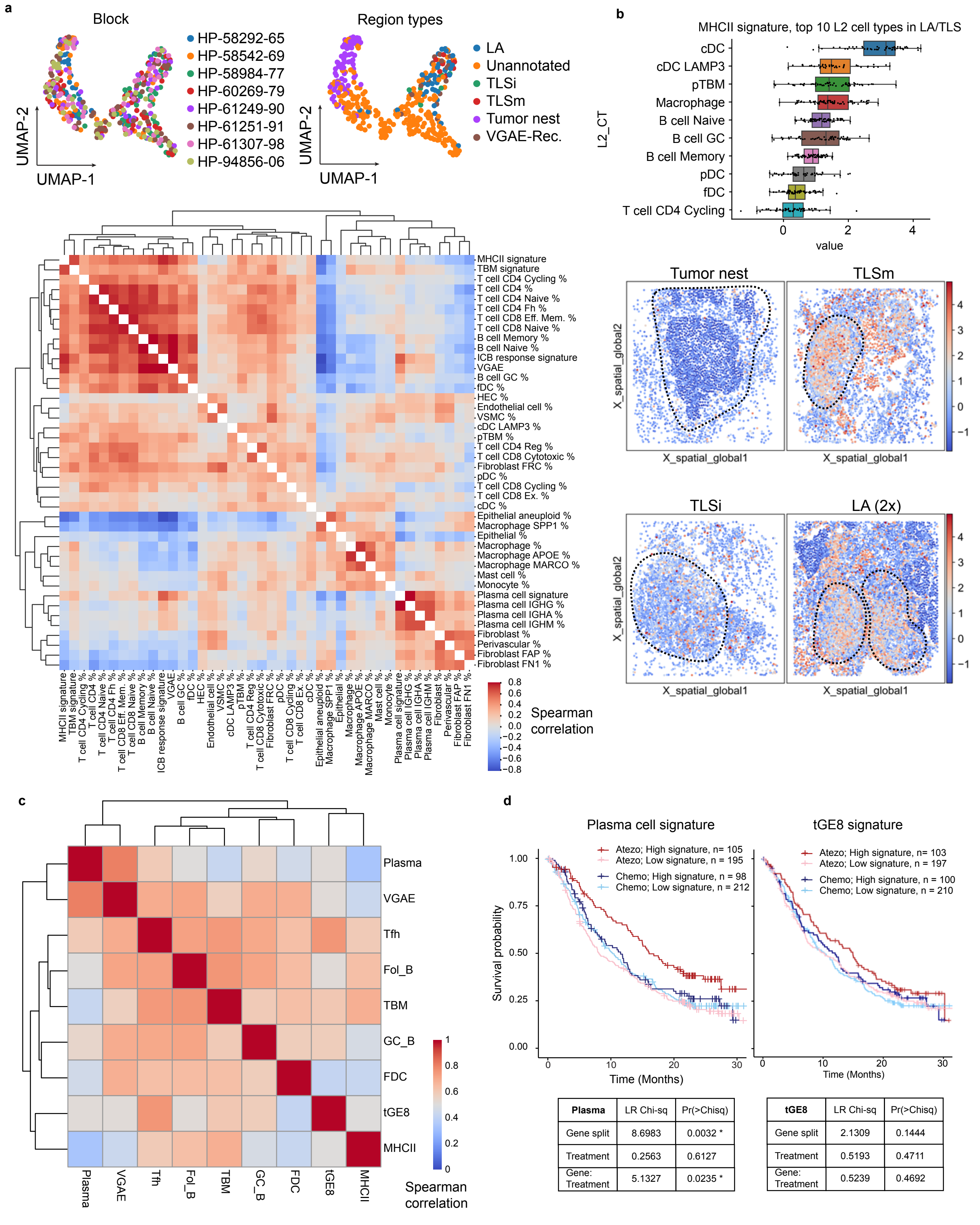


| Plasma | LR Chi-sq | Pr(>Chisq) |
|---|---|---|
| Gene split | 8.6983 | 0.0032 * |
| Treatment | 0.2563 | 0.6127 |
| Gene: Treatment | 5.1327 | 0.0235 * |

| tGE8 | LR Chi-sq | Pr(>Chisq) |
|---|---|---|
| Gene split | 2.1309 | 0.1444 |
| Treatment | 0.5193 | 0.4711 |
| Gene: Treatment | 0.5239 | 0.4692 |

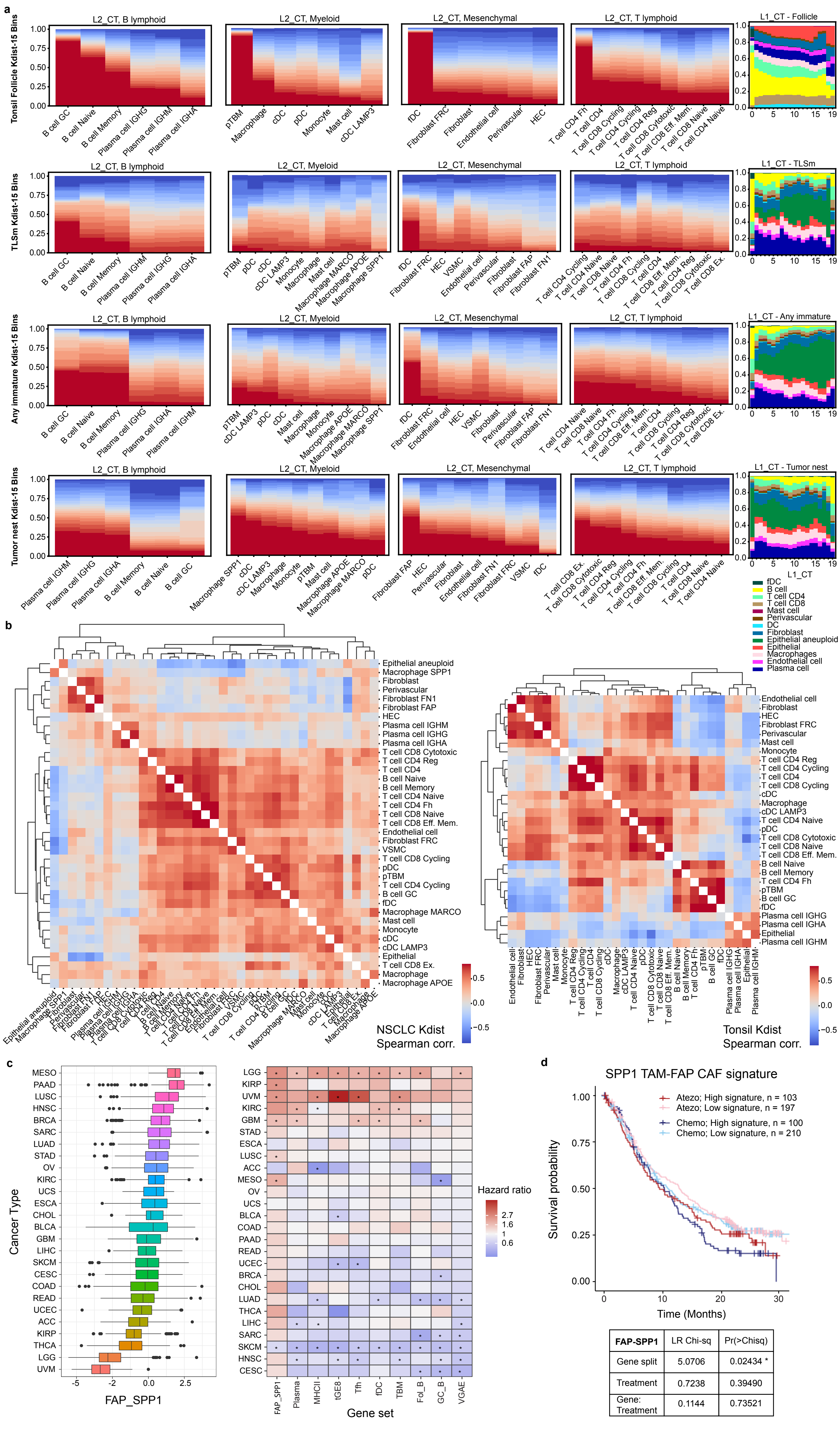
a
Tonsil Follicle Kdist-15 Bins
TLSm Kdist-15 Bins
Any immature Kdist-15 Bins
Tumor nest Kdist-15 Bins
L2_CT, B lymphoid
L2_CT, Myeloid
L2_CT, Mesenchymal
L2_CT, T lymphoid
L1_CT - Follicle
L1_CT - TLSm
L1_CT - Any immature
L1_CT - Tumor nest
L1_CT
fDC
B cell
T cell CD4
T cell CD8
Mast cell
Perivascular
DC
Fibroblast
Epithelial aneuploid
Epithelial
Macrophages
Endothelial cell
Plasma cell
b
NSCLC Kdist
Spearman corr.
Tonsil Kdist
Spearman corr.
c
Cancer Type
FAP_SPP1
Hazard ratio
Gene set
FAP_SPP1
Plasma
MHCII
tGE8
Tfh
fDC
TBM
Fol_B
GC_B
VGAE
d
SPP1 TAM-FAP CAF signature
Atezo; High signature, n = 103
Atezo; Low signature, n = 197
Chemo; High signature, n = 100
Chemo; Low signature, n = 210
Survival probability
Time (Months)
FAP-SPP1 | LR Chi-sq | Pr(>Chisq)
Gene split | 5.0706 | 0.02434 *
Treatment | 0.7238 | 0.39490
Gene: Treatment | 0.1144 | 0.73521
